# RADIF(C1orf112)-FIGNL1 Complex Regulates RAD51 Chromatin Association to Promote Viability After Replication Stress

**DOI:** 10.1101/2023.09.25.556595

**Authors:** Jessica D. Tischler, Hiroshi Tsuchida, Tommy T. Oda, Ana Park, Richard O. Adeyemi

## Abstract

Homologous recombination (HR) plays critical roles in repairing lesions that arise during DNA replication and is thus essential for viability. RAD51 plays important roles during replication and HR, however, how RAD51 is regulated downstream of nucleofilament formation and how the varied RAD51 functions are regulated is not clear. We have investigated the poorly characterized protein c1orf112/RADIF that previously scored in genome-wide screens for mediators of DNA inter-strand crosslink (ICL) repair. Upon ICL agent exposure, RADIF loss leads to marked cell death, elevated chromosomal instability, increased micronuclei formation, altered cell cycle progression and increased DNA damage signaling. RADIF is recruited to damage foci and forms a complex with FIGNL1. Both proteins have epistatic roles in ICL repair, forming a co-stable complex. Mechanistically, RADIF loss leads to increased RAD51 amounts and foci on chromatin both with or without exogenous DNA damage, defective replication fork progression and reduced HR competency. We posit that RADIF is essential for limiting RAD51 levels on chromatin in the absence of damage and for RAD51 dissociation from nucleofilaments to properly complete HR. Failure to do so leads to replication slowing and inability to complete repair.

## INTRODUCTION

Cells are constantly exposed to agents that can damage DNA (*1, 2*). These problems can be particularly inimical during cellular replication during which failure to adequately repair the lesions formed can lead to severe cellular consequences including mutations, loss of genetic information, chromosomal fusion events, and so on, all of which can lead to various diseases including cancer (*3–5*). Among the various types of lesions that can arise during replication, helix distorting crosslinks, especially inter-strand crosslinks (ICLs) in which two opposite strands are covalently ligated, can pose an absolute impediment to replication and/or transcription complex advancement and are particularly cytotoxic (*6*). ICLs can arise as byproducts of endogenous metabolic events, for example, during aldehyde and nitrous acid metabolism (*7, 8*). Several platinum-based ICL-inducing agents such as cisplatin, oxaliplatin and mitomycin C (MMC) enjoy widespread use in chemotherapy owing to the exquisite sensitivity of rapidly dividing cells to these drugs (*9, 10*).

Crosslink repair is typically coupled to replication in eukaryotes, and it involves the concerted action of various repair pathways (*6*). Among the major players are the nucleotide excision repair (NER) pathway, translesion synthesis (TLS) and HR pathways. The Fanconi anemia (FA) pathway comprises several HR genes as well as multiple genes that make up the FA core complex which, upon sensing the ICL, activates a critical complex of FANCI and FANCD2 proteins (the I-D complex) via mono-ubiquitination by the ubiquitin ligase FANCL (*11, 12*). This step is necessary for the incision events that unhook the ICL prior to NER, TLS and HR. Germline mutations affecting genes in the FA pathway cause Fanconi anemia, a rare genetic disorder characterized by bone marrow suppression, hematopoietic and growth defects and increased cancer predisposition (*13–15*).

A critical step in HR is the efficient nucleation of single-stranded DNA (ssDNA) by the recombinase RAD51 to mediate homology search, strand invasion and pairing, prior to copying of homologous strands for repair (*16, 17*). Because of its highly recombinogenic potential, this process is tightly regulated, with several factors called mediators (such as BRCA2 and the RAD51 paralogs) acting to displace the ssDNA-bound replication protein A (RPA) and promote RAD51 nucleofilament formation while others like the BLM, PARI and RECQL5 oppose and/or fine-tune the process. Downstream of filament formation, nucleofilament disassembly and subsequent DNA synthesis is also a critical but not well understood process, and only recently have important new players such as ZGRF1 and HROB (MCM8IP) been identified to promote RAD51 filament disassembly and allow postsynaptic synthesis, providing more insight into these latter steps of HR (*18–20*). All of these factors are essential for viability in response to ICL treatment.

In addition to classical HR functions of RAD51, recent work has highlighted break-repair independent functions for RAD51 during stalled fork metabolism (*21–24*). These range from promoting fork reversal, a process in which stalled forks are processed into four-way junctions to stabilize the forks, to directly protecting forks from degradation by exonucleases. Indeed, RAD51 itself is a Fanconi gene, FANCR, and certain mutants of RAD51 that are competent for HR are still quite sensitive to ICLs (*25*), demonstrating that RAD51 plays multiple roles during ICL repair. RAD51 can bind to both ssDNA and dsDNA in vitro, and although binding to dsDNA is detrimental to its HR functions, such binding has recently been shown to be critical for maintaining fork integrity (*26*). RAD51 has been shown to associate with DNA during normal replication in the absence of damage (*22*) and, while a few novel regulators of RAD51’s HR function such as FIGNL1 has been identified (*27*), there remains a need to characterize the various factors that regulate the varied repair-independent function of RAD51.

We and others previously identified c1orf112 from whole genome CRISPR screens for ICL sensitizers (*28, 29*). Here we have investigated the role of c1orf112, a previously poorly characterized factor that we now show is critical for ICL repair and RAD51 regulation. For reasons expounded upon later in the manuscript we renamed c1orf112 as ***RAD***51-regulatory ***I***nteractor of ***F***IGNL1 (RADIF). We describe c1orf112/RADIF as a novel DNA repair factor that regulates RAD51 chromatin association in the absence and persistence on nucleofilaments upon induction of exogenous DNA damage. RADIF interacts with FIGNL1 through its N-terminal region and together both proteins form a co-stable complex. Upon replication stress, loss of RADIF leads to increased genomic instability, characterized by elevated damage signaling, chromosomal aberrations, micronuclei formation, and defective HR.

## RESULTS

### C1orf112/RADIF is a novel ICL repair gene

We previously reported a genome-wide CRISPR/Cas9 knockout screen aimed at identifying novel genes required for ICL repair specifically and the replication stress response in general (*28*). This screen generated a high confidence dataset of genes not previously linked to the replication stress response. The Durocher group also reported similar cisplatin screens performed in a different cell line as part of a wider analyses of DNA repair dependencies (*29*). In order to mine high-confidence hits for follow up analysis, we selected the top-scoring 500 genes from two cisplatin screens done in RPE1 cells and two done in U2OS cells. To discover genes that function in a cell-type independent manner we looked for hits that scored in at least 3 screens. Our analysis revealed at least 70 genes, almost all of which have been linked to DNA repair (Figure 1A, Supplemental Table 1). We decided to focus on c1orf112 (RADIF), one of the uncharacterized genes on the list. RADIF scored very highly in both RPE1 cisplatin screens analyzed and in PD5 of our U2OS cisplatin screens (Figure 1A). The major isoform of RADIF encodes an 853 amino acid (aa) protein that is well conserved across vertebrates (*30*). RADIF is largely uncharacterized, however a few publications have linked high expression of RADIF to poor prognosis in various cancer types (*31, 32*).

**Fig. 1.**
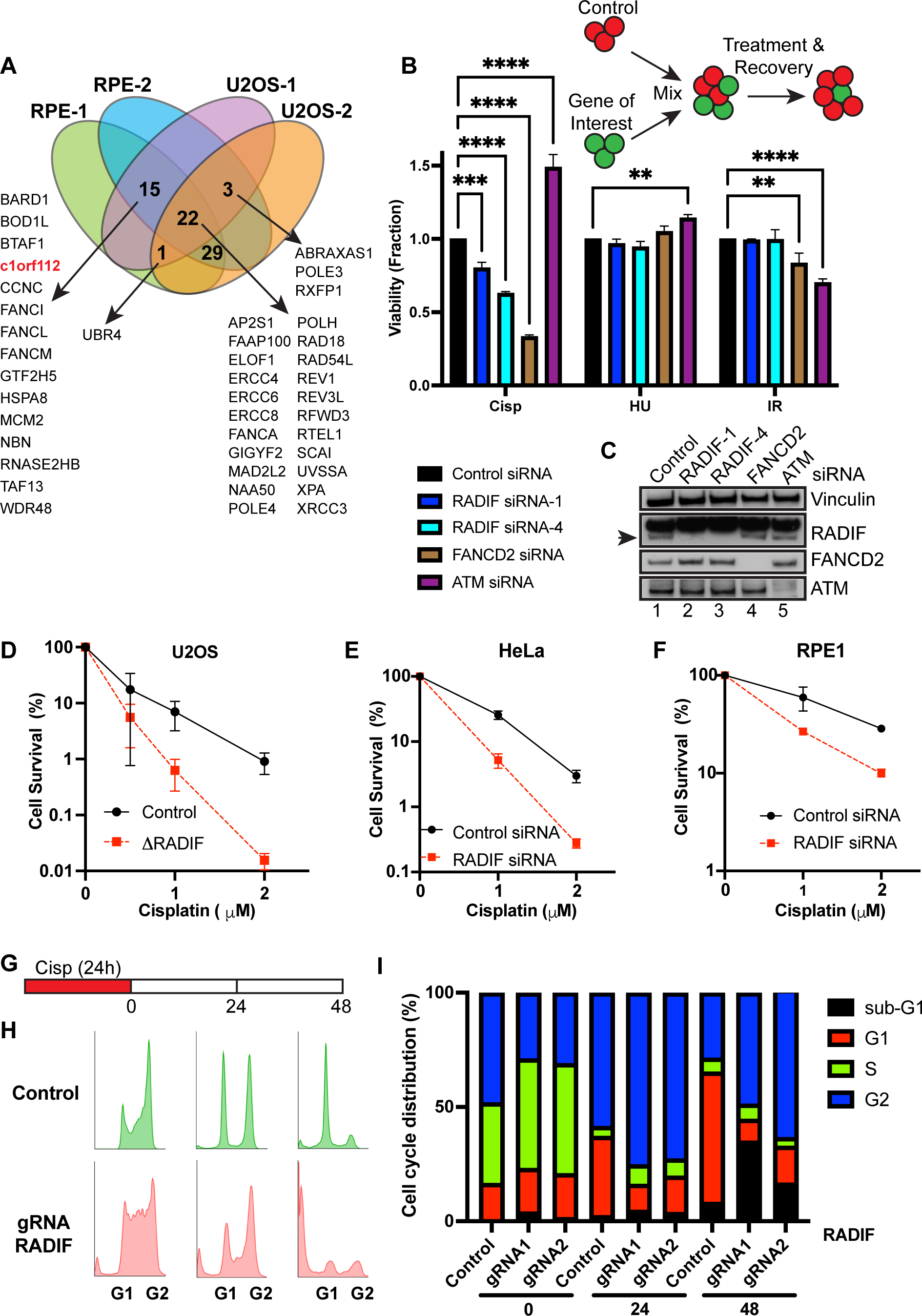
RADIF is important for ICL repair. **(A)** Schematic showing overlap of high-priority hits from published cisplatin screens. Genes that scored in at least three screens are indicated. RADIF (c1orf112) is indicated in red. All 5 overlaps are listed except for the 29 genes due to space limitations. **(B)** Top: schematic of multicolor competition assay (MCA). Middle and bottom: U2OS-GFP cells were reverse transfected and processed as described in the methods section. MCA showing survival of U2OS cells after treatment with the indicated drugs and siRNAs. Data are normalized to untreated cells. Mean and standard deviation (SD) values of two independent experiments are shown. Doses of drugs were 1 µM for cisplatin and 2 mM for hydroxyurea (HU). Ionizing radiation (IR) was performed using 9 Gy. **(C)** Western blots (WB) confirming knockdown of indicated proteins in the MCA assay. RADIF protein is indicated with an arrow below the non-specific band. **(D)** Colony survival assays (CSAs) showing survival of U2OS cells expressing control gRNA or RADIF gRNA knockout upon treatment with indicated doses of cisplatin. Mean and standard deviation (SD) of two independent experiments are shown. (**E and F)** CSAs showing survival of control or RADIF siRNA - treated HeLa cells (E) or RPE1 cells (F) upon treatment with indicated doses of cisplatin. **(G)** Schematic of cell cycle assay. Cisplatin was added at 2 μM final concentration. Cells were washed and allowed to recover for the indicated times. **(H)** Representative images of cell cycle distribution. (**I)** Cell cycle distribution in control or RADIF gRNA expressing cells at the indicated times following cisplatin treatment.

To validate a role for RADIF following replication stress, we depleted RADIF in U2OS cells using two independent validated siRNAs and performed multicolor competition assays (MCAs) (Figure 1B and C). This assay allows us to examine roles for RADIF following various genotoxic treatments. ATM (important for double strand break repair) and FANCD2 (an HR factor that is essential following various replication stress events) depletion served as controls. Whereas we failed to observe significant sensitivity following hydroxyurea treatment (which, at the doses used, stalls forks following nucleotide depletion but doesn’t damage DNA), as well as IR treatment (which causes double strand breaks and other forms of base damage), RADIF depletion significantly increased sensitivity to cisplatin treatment.

To further characterize RADIF’s role, using CRISPR-Cas9 gene editing, we generated two knockout clones in U2OS cells, (Figure S1A) and examined the knockouts for sensitivity to mitomycin C (MMC, another ICL-inducing agent) using clonogenic survival assays (CSAs). Both clones showed similar increased sensitivity to MMC upon RADIF loss (Figure S1B). We also examined cisplatin sensitivity using CSAs. Confirming a role of RADIF in these cells, we observed marked reduction in cell survival following cisplatin treatment following loss of RADIF (Figure 1D). To further define roles for RADIF in additional cell lines, we depleted RADIF using siRNAs in HeLa cells and RPE1 cells and examined cellular viability following cisplatin treatment using CSAs. Knockdown of RADIF led to significantly reduced viability in both of these cell types following treatment with the ICL-inducing agent cisplatin (Figure 1E and F). Taken together, these results from multiple cell lines demonstrate the essential function of RADIF in promoting cell viability after treatment with ICL agents.

### RADIF prevents genomic instability following replication stress

To further characterize repair roles for RADIF upon ICL agent treatment, we assayed for cell cycle alterations. RADIF loss did not cause significant changes in cell cycle distribution in the absence of damage (Fig S1C and S1D), although there was slight accumulation of cells in the G2/M population. Upon cisplatin treatment however, loss of RADIF led to marked alterations in cell cycle progression. In these experiments, cells were treated with cisplatin for a 24 h period prior to washout to allow the cells to recover for up to 2 days after drug treatment (Figure 1G). Whereas control cells showed mild accumulation of cells initially in S (0h) and later in G2 (24h after release) with almost complete recovery by day 2 after drug treatment, loss of RADIF led to marked increases in the S and G2 accumulated populations that progressed to significant apoptosis (sub G1 population) 2 days after drug treatment (Figure 1H-I).

Next, we performed metaphase spreads to examine cells for genomic instability upon RADIF loss (Figure 2A-B). RADIF knockdown led to mild increases in spontaneous chromosomal abnormalities (quantified in Figure 2B). To test the effect of ICL treatment, we employed very low MMC doses in which very little chromosomal aberrations arose in control cells. Upon MMC treatment, RADIF knockdown significantly increased the amounts of chromosomal abnormalities – breaks and gaps (Figure 2A-B). One characteristic chromosomal abnormality seen upon loss of Fanconi genes following ICL treatment is radial formation (*7, 33*). As expected, knockdown of the Fanconi gene FANCD2 led to significant increases in radial formation. However, at the low doses of MMC used in our experiments, we did not observe such increases in radial formation upon loss of RADIF (Fig S2A), suggesting that although RADIF is important for ICL repair, it may not do so as a member of the Fanconi pathway.

**Fig. 2.**
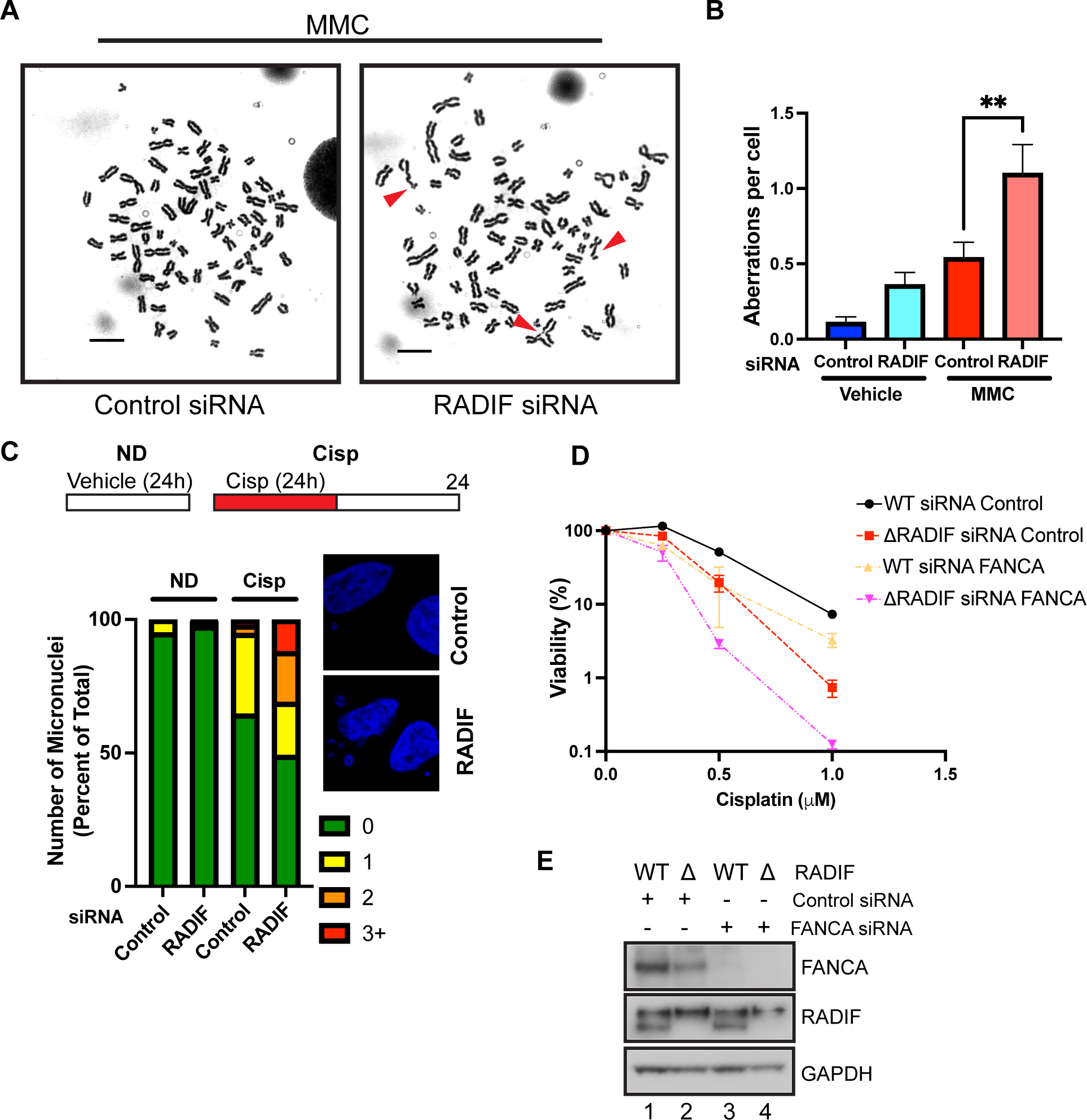
RADIF prevents genomic instability after replication stress and functions downstream of Fanconi pathway activation. **(A)** Representative images showing increased genomic instability in MMC-treated, HeLa cells following RADIF siRNA compared to control siRNA. Aberrations are shown in red arrows. Scale bar is 10 µm. **(B)** Quantification of total aberrations from experiment shown in (A). Mean ± SD of two independent experiments are shown. **(C)** U2OS cells treated with control or RADIF siRNA were exposed or not to 1 μM cisplatin for 24 h. Number of micronuclei per cell was quantified by DAPI staining 24 h after release from cisplatin treatment. ND – no drug. **(D)** U2OS cells expressing control gRNA (WT) or 1′RADIF cells were treated with control or FANCA siRNA for 48 h. Cells were then exposed to the indicated doses of cisplatin and processed for CSAs as above. Mean and SD of two independent experiments are shown. **(E)** Western blots showing knockdown of the indicated proteins from the experiment in (D).

Micronuclei formation is another hallmark of replication stress (*34*). They form as a result of lagging and/or acentric chromosomes that are not incorporated into daughter cell nuclei during cell division (*34, 35*). We treated cells with cisplatin and allowed cells to recover for 24 h before assaying for the number of micronuclei formed per cell. RADIF knockdown led to increased frequency of nuclei with higher numbers of micronuclei associated with them compared to controls (Figure 2C). There was no such increase in the absence of drug treatment. Taken together, our results show that RADIF is essential for maintaining genomic stability especially after treatment with ICL agents.

### RADIF promotes ICL repair independently of the Fanconi pathway

The striking requirement for RADIF following treatment with agents that induce DNA crosslinks prompted us to investigate whether RADIF modulates the Fanconi pathway in some way, since several Fanconi genes play prominent roles in promoting cell viability following ICL induction. During repair of DNA crosslinks, FANCM senses DNA lesions and recruits the Fanconi complex, consisting of several genes that activate FANCI-FANC2 via phosphorylation and mono-ubiquitination (*11, 36*). In order to examine whether RADIF functions as part of the Fanconi pathway, we assayed WT and ΔRADIF cells for FANCI foci. Following MMC treatment, there was increased amounts of γH2AX as well as FANCI foci (Figure S2B-C). However, we observed no significant reduction in the number of FANCI foci upon RADIF loss (Figure S2B and C). To further determine whether RADIF modulates Fanconi pathway activation, we performed immunoblotting to directly assay for mono-ubiquitination of FANCD2, seen as a slight reduction in protein mobility. Cisplatin treatment led to increased mono-ubiquitination of FANCD2, with no apparent reduction in this process upon RADIF depletion (Figure S2D, compare lanes 5 and 6 to lane 4).

To further characterize RADIF’s potential interaction with the Fanconi pathway, we depleted FANCA in WT or ΔRADIF cells. FANCA is core member of the complex that mono-ubiquitinates and activates FANCI-D2, thus FANCA depleted cells are very sensitive to treatment with ICL agents. Near complete FANCA depletion was confirmed by western blotting (Figure 2E). ΔRADIF cells showed similar sensitivity to ICLs as compared FANCA knockdown cells, with ΔRADIF cells showing slightly higher sensitivities as the dose of drug increased (Figure 2D). Importantly, co-depletion of FANCA and RADIF led to increased reduction in viability compared to depletion of either protein alone (Figure 2D-E). Taken together, while RADIF plays vital roles in promoting cellular viability after ICL treatment, it appears to do so in a manner that is distinct from but additive to the roles played by the FA pathway.

### RADIF is recruited to DNA damage sites and promotes repair following DNA damage

To visualize RADIF localization following DNA damage we performed immunofluorescence (IF) assays using antibodies against epitope–tagged RADIF following cisplatin treatment. RADIF formed multiple foci upon cisplatin treatment (Figure 3A and B). Similar results were seen in both U2OS and HeLa cells (Fig S3A). There were some vehicle-treated cells that showed RADIF foci formation (Figure 3B). RADIF foci only partially colocalized with γH2AX (Figure 3A-B), and in many cases was situated right adjacent to γH2AX foci. Knockdown of FANCD2 almost completely abolished foci formation by RADIF after cisplatin treatment (Figure 3C-D), despite little to no colocalization with FANCD2 (Figure S3B), suggesting that RADIF acts downstream of FANC-ID foci formation but upstream of CtIP, whose knockdown did not prevent RADIF foci formation (Figure 3C-D).

**Fig. 3.**
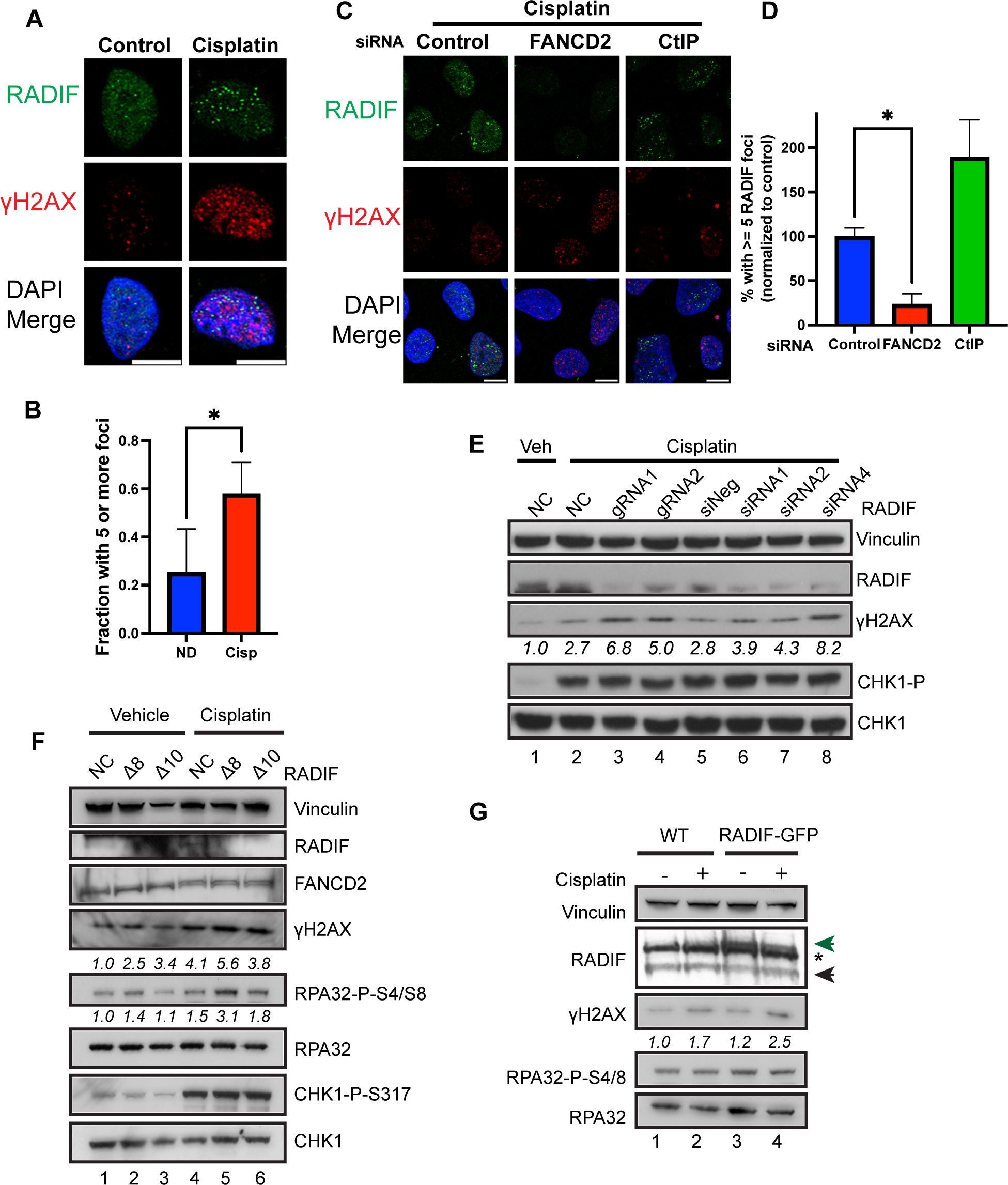
RADIF is recruited to damage foci and limits DNA damage signaling after ICL treatment. **(A)** IF images showing recruitment of GFP-RADIF to foci that partially colocalize with γH2AX after treatment with 0.5 μM cisplatin for 16 h. Cells were pre-extracted to remove soluble protein. **(B)** Quantification of the experiment in (A). Mean ± SD of four independent experiments are plotted. **(C-D)** U2OS cells were transfected with the indicated siRNAs for 48 h prior to treatment with 0.5 μM cisplatin for 16 h. IF images show reduced recruitment of GFP-RADIF to foci following loss of FANCD2. Quantified in (D). Mean ± SD of two independent experiments are plotted. **(E)** U2OS cells expressing control or two independent gRNAs to RADIF (lane 1-4), or U2OS cells 48 h after reverse transfection with control siRNAs or three independent siRNAs to RADIF (lane 5-8) were treated with vehicle or 1.5 μM cisplatin for 16 h prior to western blotting for the indicated proteins. Blots show increased γH2AX levels following knockdown of RADIF using siRNAs and gRNAs. γH2AX numbers are normalized to vinculin loading control. **(F)** Control gRNA expressing or two independent 1′RADIF U2OS clones were exposed to vehicle or 2 μM cisplatin for 18 h prior to blotting for the indicated proteins. Numbers are normalized to vinculin loading control (γH2AX) or total RPA (pRPA32). **(G)** Control U2OS or U2OS cells stably expressing near-endogenous levels of GFP-RADIF (were exposed to vehicle or 2.5 μM cisplatin for 18 h prior to blotting for the indicated proteins. γH2AX numbers are normalized to vinculin loading control. * - non-specific band. Black arrowhead – endogenous RADIF protein. Green arrowhead – GFP-RADIF.

Next, we examined for any differences in DNA damage signaling upon RADIF loss. During replication stress, RPA-coated ssDNA at stalled replication fork junctions trigger activation of ATR (*37–39*). This initiates a signaling cascade in which several proteins ranging from repair genes to checkpoint modulator are activated. While RADIF knockdown had no significant effect on the level of phosphorylation of the checkpoint protein Chk1 (Figure 3E), RADIF knockdown consistently led to increased γH2AX and RPA phosphorylation (Figure 3E and 3F) in the presence of DNA damage, as well as to mild increases in phosphorylation of both proteins in the absence of exogenous damage (Figure 3F). This was seen both following gRNA and siRNA depletion of RADIF (Figure 3E), with multiple gRNAs and siRNAs showing effects correlating to the level of RADIF knockdown.

RADIF has been observed to be overexpressed in multiple cancer cell types (*31, 32*). To further characterize RADIF’s role on genomic integrity, we generated cells that stably expressed near-endogenous levels of GFP-tagged RADIF (thus near 2-fold total expression). Such modest RADIF overexpression also showed similar increases in the levels of γH2AX (Figure 3G) both in the presence (Figure 3G, compare lanes 2 and 4) and absence of exogenous DNA damaging agents. Taken together, the increased damage signaling observed both when RADIF was lost or overexpressed showed that alterations in RADIF levels was highly correlated with dysregulated DNA damage signaling, suggesting a need for tight regulation of cellular RADIF levels to prevent genomic instability.

### RADIF interacts with FIGNL1

BioGrid interactomes obtained from published mass spectrometry datasets suggest interactions between RADIF and FIGNL1 (*40*). FIGNL1 was originally identified in a proteomics dataset for interactors of RAD51 (*27*). Examining the reported IP mass spec dataset for interactors of FIGNL1 showed multiple peptides to c1orf112/RADIF. Indeed, the *Arabidopsis* ortholog of RADIF, named FLIP, shows interaction between the two proteins (*41*). To determine whether both mammalian proteins interact, we expressed GFP-tagged RADIF and FLAG-tagged FIGNL1 in 293 cells. Using co-immunoprecipitation (coIP) experiments, FIGNL1 was able to pull down tagged RADIF, validating their interaction (Figure 4A). Likewise, expression of tagged-RADIF pulled down the endogenous FIGNL1 protein (Figure 4B). Notably, the interaction was constitutive, and did not require treatment with DNA damaging agents. To determine whether DNA damage signaling enhanced the interaction between the two proteins, we performed the coIPs in the presence or absence of cisplatin. To our surprise, cisplatin treatment significantly reduced the interaction between RADIF and FIGNL1 (Figure 4C, compare lanes 3 and 4).

**Fig. 4.**
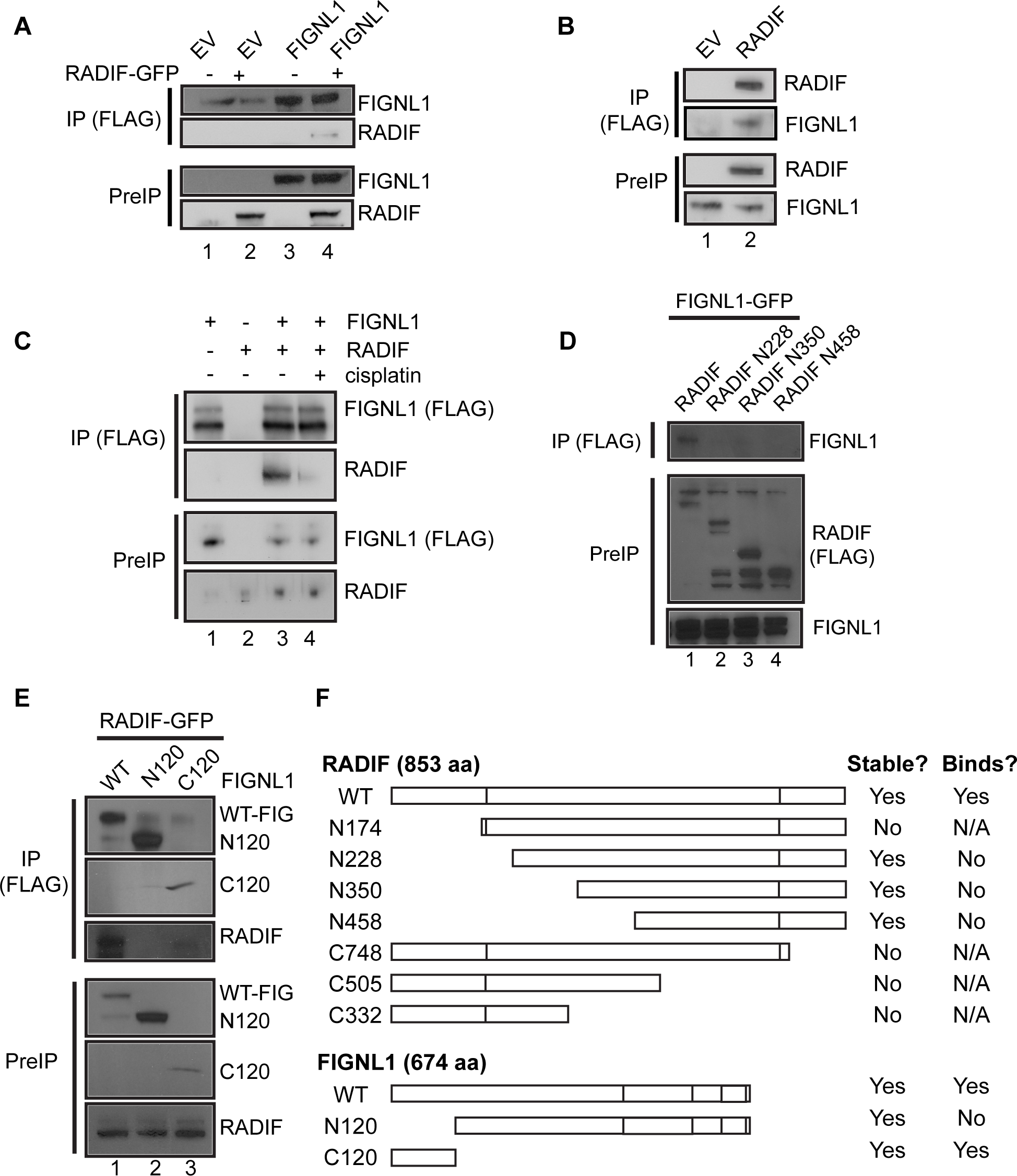
RADIF forms a complex with FIGNL1. **(A)** 293T cells were transfected with empty vector (EV), FLAG-FIGNL1 or GFP-RADIF as indicated for 48 hrs. Blots show association of FIGNL1 and RADIF. **(B)** 293T cells were transfected with EV or FLAG-RADIF for 48 h. Western blots show pull down of endogenous FIGNL1 following coIPs. **(C)** 293T cells were transfected with EV, FLAG-FIGNL1 or GFP-RADIF as indicated for 48 h. Cells were then treated with 2 μM cisplatin for 24 h prior to coIP. Blots show reduced association of FIGNL1 and RADIF upon cisplatin treatment. **(D)** 293T cells were transfected with FIGNL1-GFP together with either full-length RADIF-FLAG or the indicated truncation mutants for 48 h prior to coIP and blotting for the indicated proteins. **(E)** 293T cells were transfected with RADIF-GFP and either full-length FIGNL1-FLAG or the indicated truncation mutants for 48 h prior to coIP and blotting for the indicated proteins. **(F)** Schematic showing the FIGNL1 and RADIF truncations and the coIP results. Mutants are drawn to scale.

Next, to map the domain that mediates binding between RADIF and FIGNL1, we generated a series of N and C-terminal truncation mutants of RADIF. A few of these mutants were unstable and could not be expressed (summarized in Figure 4F). However, our experiments revealed that a mutant lacking the N-terminal 227 aa was severely compromised in its ability to bind to FIGNL1 even though it expressed better than the wildtype in transient transfections (Figure 4D), suggesting that binding between RADIF and FIGNL1 requires this N-terminal region. Further deletions beyond the first 350 aa of the protein completely abolished the interaction (Figure 4D). Although the N-terminal 227 aa was required, a truncation mutant expressing only the N-terminal portion of the protein was unstable, precluding us from determining whether the N-terminal region was sufficient to bind to FIGNL1. (These C-terminal truncations were tagged with an NLS, since the predicted NLS of RADIF is located at the C-terminus of the protein ruling out mislocalization as the reason for the instability of the protein).

To map the interaction region of RADIF on FIGNL1, we started by generating two previously reported mutants that divide the FIGNL1 protein into an N-terminal 120aa region and a large truncation lacking the N-terminus (Figure 4F). This latter mutant was previously shown to be defective in FIGNL1’s HR functions upon DNA damage. In addition to 3X-FLAG tags, the N-terminal region was tagged with an SV40-dervied NLS, since this mutant was previously shown to localize to the cytoplasm when expressed unlike the full-length protein and the C120 mutant that both localize to the nucleus. Expressing the three proteins in 293 cells, coIP experiments revealed a striking loss of interaction upon deletion of the N-terminal region (Figure 4E). Importantly, expression of only this region was sufficient to pull down RADIF (Figure 4F) although at reduced levels, as this mutant was expressed at much lower levels than the full-length protein. Taken together, we conclude that both FIGNL1 and RADIF bind to each other using their respective N-terminal regions.

### RADIF / FIGNL1 function in the same pathway to promote viability after ICL treatment

Both RADIF and FIGNL1 scored in genome-wide cisplatin screens, and DepMap analysis for dependents of RADIF showed a high correlation between loss of RADIF and loss of FIGNL1 (*42*), suggesting that both proteins not only interact but likely functioned in the same pathway. Indeed, the N-terminal deficient mutant of FIGNL1 was unable to rescue HR defects upon FIGNL1 depletion and was not recruited to DNA damage sites, suggesting that FIGNL1 loss may be epistatic with RADIF loss (*27*). To test this, we depleted FIGNL1 using siRNAs in WT vs RADIF knockout cells and examined sensitivity to cisplatin by CSAs. Loss of FIGNL1 led to increased sensitivity to cisplatin that was almost as high as RADIF knockouts (Figure 5A). Importantly, unlike what was seen with FANCA depletion (Figure 2G-H), depletion of FIGNL1 in ′RADIF cells was no different than control siRNA treatment, suggesting that both proteins function in the same pathway in mediating survival after replication stress (Figure 5A).

**Fig. 5.**
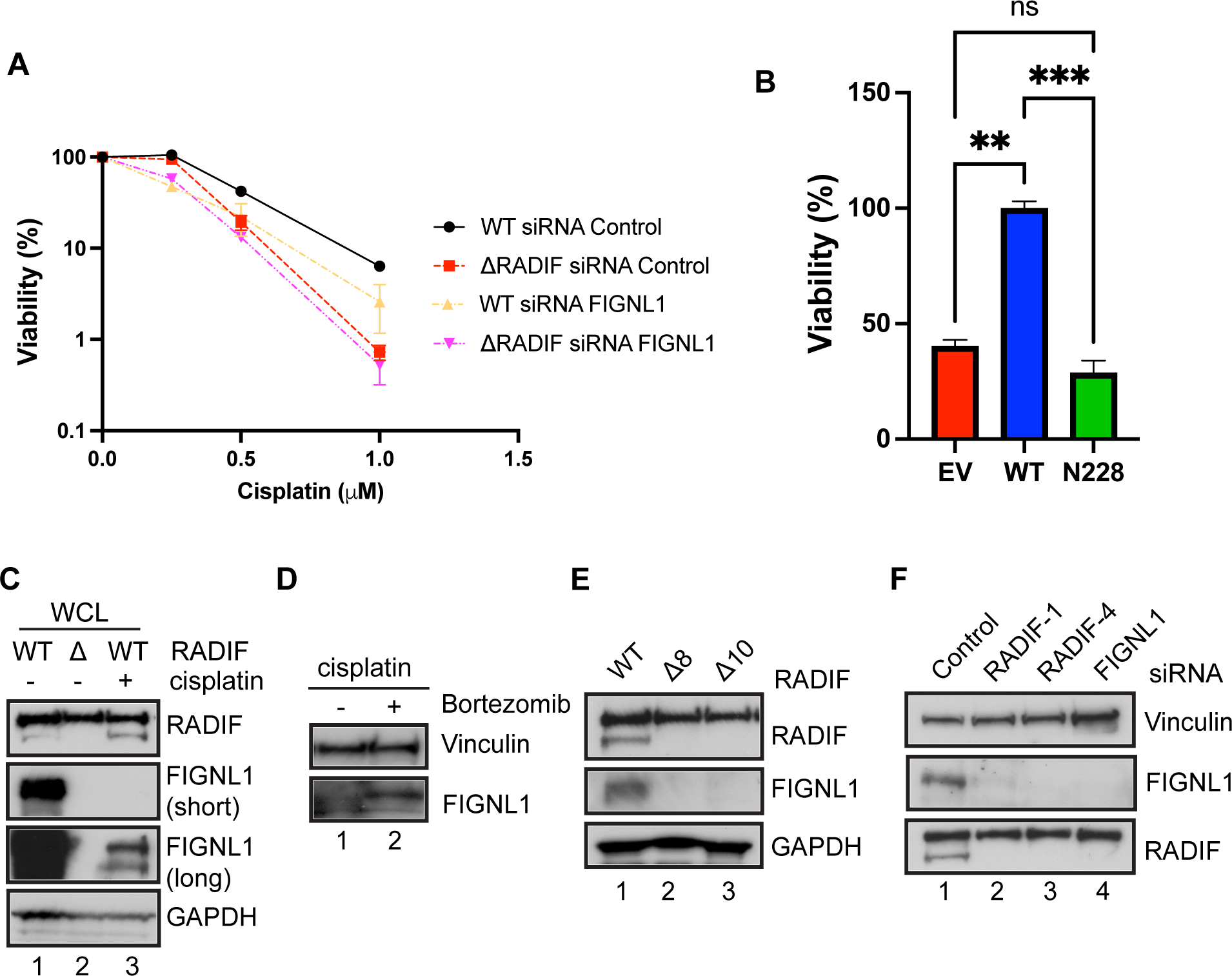
RADIF and FIGNL1 function in the same pathway and form a co-stable complex. **(A)** U2OS cells expressing control gRNA (WT) or RADIF null cells were treated with control or FIGNL1 siRNA for 48 h. Cells were then exposed to the indicated doses of cisplatin and processed for CSAs as above. Mean and SD of two independent experiments are shown. **(B)** ΔRADIF U2OS cells stably expressing EV, full length RADIF-FLAG (WT) or the N228 RADIF truncation mutant were exposed to 1 μM cisplatin for 24 h. Cells were then processed for CSAs as above. Mean and SD of two independent experiments are shown. **(C)** U2OS cells expressing control gRNA (WT) or RADIF null were treated or not with 1.5 μM cisplatin for 18h as indicated, prior to blotting with the indicated antibodies. WCL – whole cell lysates. **(D)** WT-U2OS cells were treated with 2.5 μM cisplatin for 14 h prior to addition of 10 μM bortezomib or vehicle for 4 h. Cells were processed for WB against the indicated proteins. **(E)** WCLs from U2OS cells expressing control gRNA (WT) or two independent ΔRADIF clones were assayed for the indicated proteins by WB. **(F)** U2OS cells were reverse transfected with the indicated siRNAs for 48 h. WCLs were prepared and WB were performed against the indicated proteins.

To further determine whether both proteins functioned together, we reasoned that RADIF mutants that failed to interact with FIGNL1 might be defective in rescuing RADIF knockout cells compared to the full-length RADIF. To test this, we complemented ′RADIF cells with vector, WT-RADIF as well as a RADIF mutant lacking the N-terminal 227aa, a mutant that we previously showed failed to interact with FIGNL1 (Figure 4D). Consistent with both proteins functioning in the same pathway to mediate cisplatin resistance, the FIGNL1 interaction-defective mutant of RADIF was unable to mediate resistance to cisplatin compared to the WT (Figure 5B). Taken together, our data suggest that RADIF and FIGNL1 function in the same pathway to mediate cellular viability after exposure to replication stress agents like cisplatin.

### RADIF and FIGNL1 form a co-stable complex

Despite both proteins appearing to function in the same pathway upon ICL reagent treatment, our coIP experiments unexpectedly showed reduced interaction between FIGNL1 and RADIF upon cisplatin treatment (Figure 4C). To understand why this was the case, we examined steady state levels of FIGNL1 and RADIF in the presence and absence of drug treatment. To our surprise, we observed a striking loss of expression of the FIGNL1 protein in the RADIF null cells (Figure 5C, compare lane 1 and 2). This was not an artifact peculiar to this knockout (KO) clone as two different KO clones showed similar losses in FIGNL1 expression (Fig 4E, compare lanes 2 and 3 to lane 1). This suggested that RADIF stabilizes the FIGNL1 protein. Since gRNA cloning takes several weeks and could lead to compensatory genomic alterations in other genes, and to determine whether short term depletion of RADIF affected FIGNL1 levels, we depleted RADIF with two different siRNAs. FIGNL1 depletion served as a control. Acute depletion of RADIF using two different siRNAs led to reduced levels of FIGNL1. To our surprise, siRNA depletion of FIGNL1 also led to loss of RADIF, demonstrating that not only does RADIF stabilize FIGNL1, but both proteins appear to form a mutual, co-stable complex.

### FIGNL1 is degraded upon ICL agent treatment

In addition to FIGNL1 loss in the absence of RADIF, we also observed reduced levels of FIGNL1 upon cisplatin treatment (Figure 5C, compare lanes 1 and 3), albeit less severely than seen following RADIF loss. This was not due to downregulation of RADIF, as RADIF levels did not change upon cisplatin treatment (Figure 5C). Unlike the reduced FIGNL1 levels seen in the absence of RADIF (which could not be reversed by proteasome inhibition (Fig S4A)), the damage-induced reduction in FIGNL1 was proteasome dependent and could be reversed by short term treatment with the proteasome inhibitor Bortezomib (Figure 5D). Taken together, FIGNL1 appears to be downregulated in a proteasome-independent manner upon RADIF loss, but targeted for proteasomal degradation upon cisplatin treatment when RADIF is present. These results likely explain the reduced observed interaction between transiently transfected RADIF and FIGNL1 upon DNA damage.

### RADIF interacts with and regulates RAD51 association with chromatin

FIGNL1 interacts with RAD51 through an FXXA domain that is present in several RAD51 binding proteins (*27, 43*). Since RADIF interacts with FIGNL1, and both proteins form a complex, we tested whether RADIF could also bind to RAD51. CoIP experiments using HA-tagged RAD51 and FLAG-tagged RADIF confirmed that RADIF interacts with RAD51 (Figure 6A). The amount of RAD51 pulled down with RADIF was lower than with FIGNL1 coIP, however, RADIF was expressed at much lower levels (Figure 6A, PreIP, compare lane 2 and 3).

**Fig. 6.**
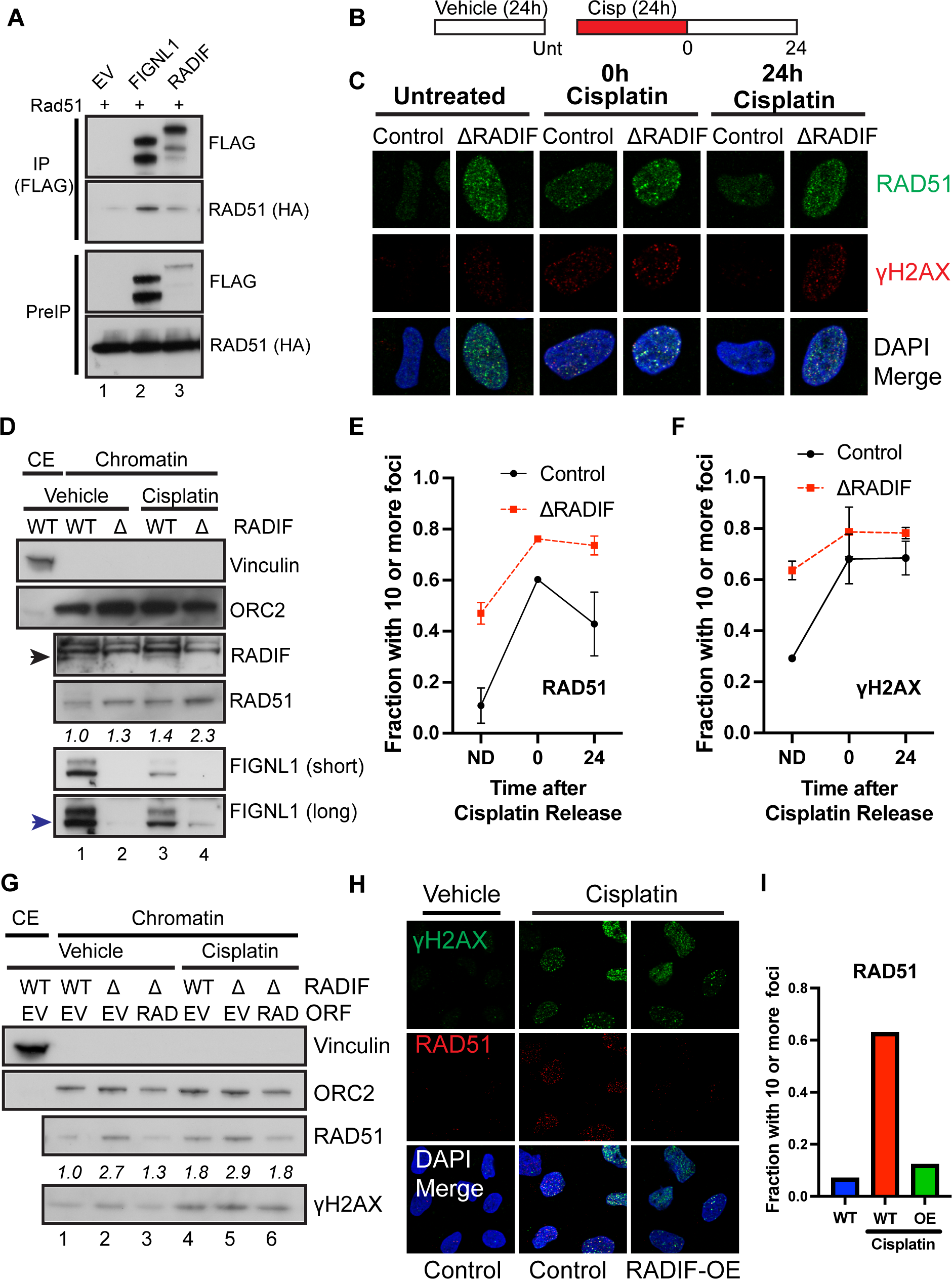
RADIF-FIGNL1 limit RAD51 chromatin association in absence of DNA damage and nucleofilament disassembly. **(A)** 293T cells were transfected with RAD51-HA and either empty vector (EV), or the indicated 3X-FLAG-tagged proteins for 48h prior to coIP using FLAG beads. **(B)** Schematic of the experiment in (C, E, F). Cisplatin was added at 0.5 μM. **(C)** Control gRNA expressing and 1′RADIF U2OS cells were processed as shown in (B). Representative IF images show RAD51 and γH2AX foci. **(D)** Control gRNA-expressing (WT) or 1′RADIF U2OS cells were treated with vehicle or 2.5 μM cisplatin for 18 h then fractionated prior to blotting for the indicated proteins. CE – cytoplasmic extract. Black arrowhead – RADIF. Blue arrowhead – FIGNL1. **(E & F)** Quantification of the experiment in (B & C). Around 100 or more nuclei were counted per condition per replicate. Data represent mean and SD of two independent replicates. ND – no drug. **(G)** Control gRNA-expressing (WT) or 1′RADIF U2OS cells stably expressing empty vector (EV) or GFP-RADIF (RAD) were treated with vehicle or 1.5 μM cisplatin for 18 h. Samples were processed for chromatin fraction prior to blotting for the indicated proteins. CE – cytoplasmic extract. RAD51 numbers are quantified related to ORC2 loading control. **(H)** Control or RADIF-GFP overexpressing U2OS cells were treated or not with 0.5 μM cisplatin for 18 hr as indicated. Representative IF images show RAD51 and γH2AX foci. **(I)** Quantification of (H).

During repair, RAD51 loading onto ssDNA leads to detectable foci formation. To determine whether RADIF modulates this process of HR, we sought to determine whether RADIF regulated RAD51 foci formation. 24h after cisplatin treatment (0h after release), RADIF depletion did not lead to reduction in RAD51 foci formation (Figure 6B, C and E). Instead, upon RADIF loss, we observed increases in the percentage of cells with greater than 10 RAD51 foci as well as increased in RAD51 nuclear staining (Figure 6C,E). This was seen both in τιRADIF cells (Figure 6C and E) as well as upon siRNA depletion of RADIF (Figure S5A-B). Due to spontaneous DNA damage formation in cells undergoing replication stress, a few foci of RAD51 are often observed in the absence of exogenous DNA damage. Strikingly, we observed significant increases in the number of RAD51 foci upon RADIF loss in untreated cells (Figure 6C and E).

To further characterize RADIF effects on RAD51 foci formation, cells treated with cisplatin for 24 h were allowed to recover for one or more days after drug washout. Interestingly, loss of RADIF led to persistence of RAD51 foci following cisplatin treatment (Figure 6C and E). This was also seen after siRNA treatment to deplete RADIF (Figure S5A-B) and RAD51 persisted at higher amounts in ′RADIF cells even 3 days after release from treatment (Figure S5C-D). These results suggest that RADIF regulates dissociation of RAD51 nucleofilaments upon DNA damage and provide a possible mechanism for DNA repair defects seen upon RADIF loss after ICL agent treatment.

As an alternative approach to examine the effects of RADIF on RAD51, we performed chromatin fractionation experiments in WT and ΔRADIF cells. We consistently observed increases in the amounts of RAD51 in chromatin fractions whenever RADIF was absent (Figure 6D, S5E). This was not due to reduction in RAD51 expression, as RADIF loss did not lead to downregulation of RAD51 levels in whole cell lysates (Figure S5E). Such increase in chromatin associated RAD51 after DNA damage was also observed after siRNA depletion of either RADIF or FIGNL1 (Figure S5F) and could be rescued by expressing tagged RADIF (Figure 6G, compare lanes 2&3 or lanes 5&6).

RADIF’s binding partner, FIGNL1, can dissociate RAD51 from nucleofilaments in vitro (*44*).To determine whether RADIF actively promotes RAD51 removal from foci, and to rule out whether the increased RAD51 foci seen in the absence of RADIF was not as a result of increased break formation (Figure 2), we took advantage of cells modestly over-expressing GFP-tagged RADIF (Figure S6G). We reasoned that RADIF overexpression should prevent RAD51 foci formation regardless of γH2AX status. Indeed, whereas both RADIF knockout and RADIF-overexpressing cells (RADIF-OE) had increased γH2AX staining (Figure S5H, see also Figure 2), RADIF-OE cells failed to form RAD51 foci (Figure S5H-I) in the absence of exogenous DNA damage. To further examine this, we treated control cells with cisplatin at doses that induced robust γH2AX and RAD51 foci formation (see Figure 6C, E, F). Compared to WT cells, RADIF-OE cells failed to form RAD51 foci even upon cisplatin treatment (Figure 6H-I) demonstrating either defective nucleofilament formation or, more likely, dissociation of RAD51 from nucleofilaments. Taken together, our data identifies RADIF and FIGNL1 as critical regulators of RAD51 chromatin association and nucleofilament disassembly upon DNA damage.

### RADIF promotes replication fork integrity

RAD51 association with DNA can drive fork reversal which might be expected to slow replication fork progression (*24*). Our observation of increased RAD51 association with DNA upon RADIF loss even in cells without exogenous DNA damage prompted us to determine whether RADIF loss affected replication fork progression. We sequentially pulsed cells with the DNA analogs CldU and IdU and, using single-molecule DNA fiber assays, examined fiber lengths of the second signal upon RADIF depletion. As a positive control we treated cells with the DNA polymerase alpha inhibitor CD437, whose inhibition was recently shown to slow both leading and lagging strand progression (*45*). We found that RADIF loss led to significant reduction in the length of the IdU signal, consistent with slowing of replication fork progression in the absence of RADIF (Figure 7A and B). Similar results were seen when RADIF was depleted by siRNA as was seen in ΔRADIF cells. Unlike CD437 treatment, we did not observe significant increases in fork asymmetry, suggesting that RADIF loss might not in itself lead to uncoupling of the leading and lagging strands (Figure S6A-D).

**Fig. 7.**
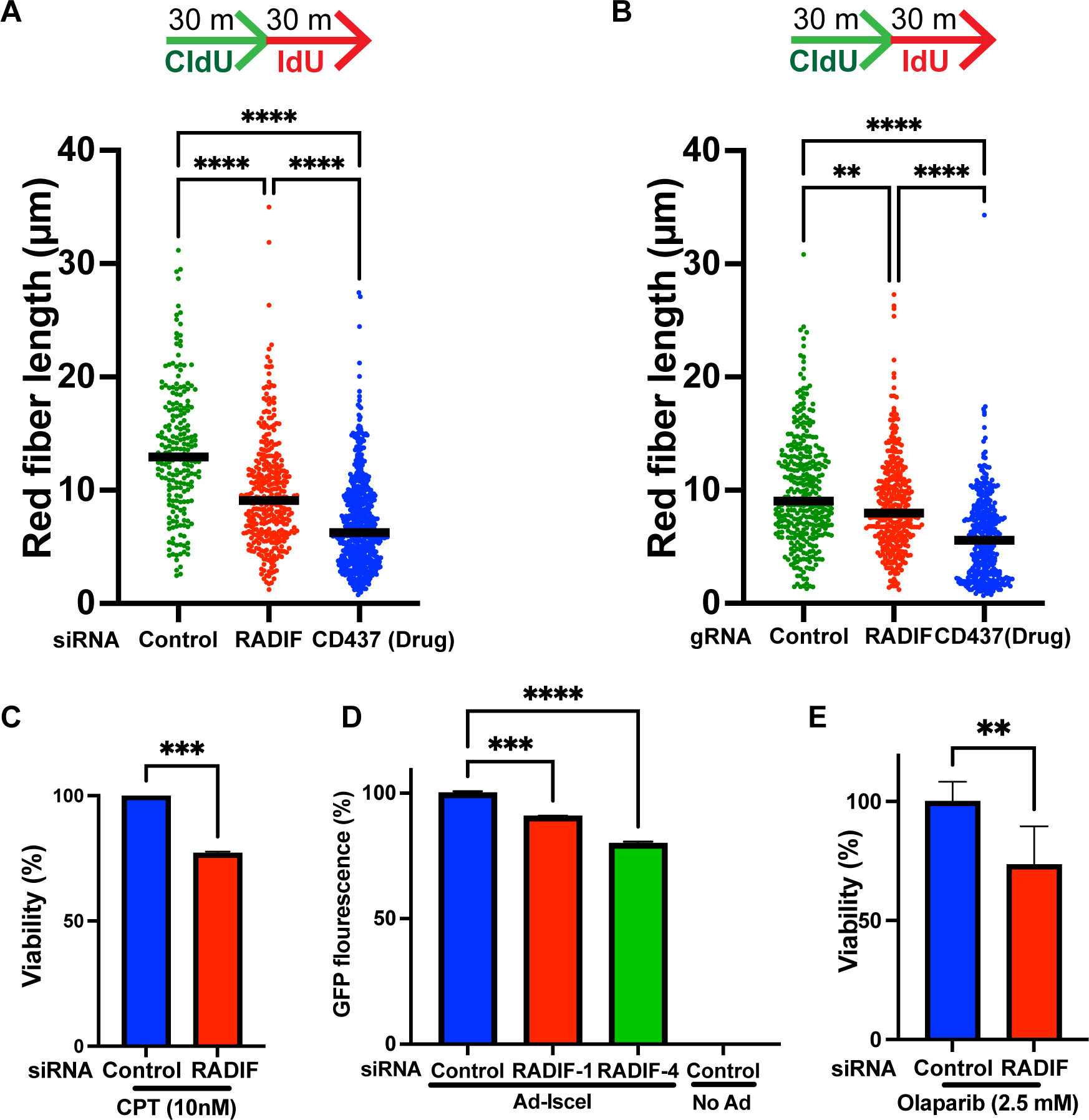
RADIF promotes replication fork progression and HR. **(A)** Schematic of pulse-labeling DNA strands, and red fiber length of two-color DNA fibers. DNA fibers were from U2OS cells transfected with siRNAs as indicated or treated with CD437. **(B)** Schematic of pulse-labeling DNA strands, and red fiber length of two-color DNA fibers. DNA fibers were from U2OS cells expressing control gRNA, RADIF gRNA or control gRNA with CD437 treatment. **(C)** U2OS-GFP cells were transfected with control or RADIF siRNA. MCA showing survival of U2OS-GFP cells after treatment with the indicated dose of camptothecin (CPT). Data is normalized to viability of control siRNA. Mean and SD of three independent experiments are shown. **(D)** U2OS DR-GFP reporter cells were reverse transfected with the indicated siRNAs for 48h and then infected with recombinant Adenovirus expressing I-SceI to cause a DSB within the reporter. 48h later, GFP-positive cells, indicating HR efficiency, were detected by flow cytometry. Data is normalized to viability of control siRNA. Mean and SD of two independent experiments are shown. **(E)** SUM149 cells were reversed transfected with control or RADIF siRNA for 48h prior to treatment with the indicated dose of Olaparib for 24h. After 2-3 days cellular viability was assayed by CellTiterGlo. Data is normalized to viability of control siRNA. Mean and SD of two independent experiments are shown.

### RADIF is important for proper homologous recombination and sensitizes BRCA mutant cells to PARP inhibitor treatment

Having observed interaction between RADIF and RAD51, and altered RAD51 dynamics in the absence of RADIF, we sought to determine whether RADIF loss regulated HR. Accumulation of RAD51 on chromatin in the absence of RADIF might suggest anti-recombinase properties for RADIF, however, the requirement for RADIF following cisplatin treatment (Figure 1) suggests that RADIF is important for HR and that increased RAD51 was likely observed due to a failure to complete HR. FIGNL1, which phenocopies RADIF loss with respect to RAD51 chromatin levels, has also been linked to reduced HR competency upon double stranded break (DSB) formation. Consistent with a requirement for RADIF for proper HR, MCA assays showed that RADIF knockdown led to reduced viability after camptothecin (CPT) treatment (Figure 7C), which causes replication-associated breaks that are repaired by HR (*46*).

To directly examine HR competency at DSBs in the absence of RADIF, we took advantage of U2OS cells expressing a previously reported modified GFP reporter that fluoresces green when an I-SceI enzyme-induced DSB is repaired (*47*). Using this assay, depletion of RADIF using two independent siRNAs led to significant HR reduction in a manner that correlated with the potency of the siRNAs to deplete RADIF (Figure 7D). To further characterize a role for RADIF in regulating HR, we examined sensitivity to PARP inhibitor treatment. PARP inhibitors are synthetic lethal with loss of HR factors and have recently been approved to treat cancers with BRCA1/2 mutations (*48–50*). To study modulation of PARP inhibitor sensitivity by RADIF loss, we made use of SUM149 cells breast cancer cells, which have mutations in the BRCA1 gene (*51*). RADIF knockdown sensitized SUM149 cells to PARP inhibitor treatment (Figure 7E). Taken together, our results demonstrate an HR function for RADIF and suggest that RADIF inhibition might be a viable way to enhance sensitivity of BRCA1 mutant cells to PARP inhibitor treatment.

## DISCUSSION

Here we have identified a novel protein, c1orf112, which we renamed RADIF, that plays vital roles in ICL repair and homologous recombination. Tight regulation of RAD51 levels is critical for proper recombination control. Our work identifies RADIF as a novel regulator of RAD51, determining its chromatin loading and foci formation both in the presence and absence of DNA damage. We envisage at least two roles for RADIF. During normal replication, RADIF associates with chromatin where it regulates RAD51 levels, allowing adequate fork progression. However, following DNA damage, the levels of and interaction between RADIF and its effector FIGNL1 is reduced, allowing increased chromatin association of RAD51, and mediating RAD51 dissociation from nucleofilaments upon completion of HR.

Although ΔRADIF cells showed exquisite sensitivity to ICL agents in particular, RADIF does not appear to be a Fanconi pathway gene. Indeed, FANC-ID activation and foci formation occurred normally in ΔRADIF cells, and although RADIF limits genomic instability after ICL treatment, we did not observe significant increases in characteristic radial formation often observed following loss of Fanconi genes. Recently, a Fanconi-independent and NEIL3-dependent sub-pathway of ICL repair was identified (*36, 52*), whether RADIF functions downstream of both pathways is not clear. However, depletion of either FANCA or RADIF led to similar reduction in viability after ICLs, which was further increased upon loss of both proteins, pointing to RADIF acting in parallel to or, more likely, downstream of FANC-ID foci activation as might be expected since RAD51 nucleofilament formation and HR occurs late during ICL repair. Indeed, FANCD2 depletion prevented RADIF foci formation upon cisplatin treatment.

RADIF’s role is closely tied to FIGNL1, with which it forms a co-stable complex. DepMap analysis revealed high correlation between loss of either gene, suggesting similar functions. We confirmed that both proteins interact and mapped the regions necessary for binding. FIGNL1 appears to bind to RADIF through its N-terminal 120 aa domain, which was both required and sufficient for binding to RADIF despite the reduced expression of this mutant compared to the WT. Interestingly, FIGNL1 mutants deficient in the N terminal 120 aa were previously shown to be defective in recruitment to damage sites and in promoting HR (*27*). However, although this was consistent with the idea that RADIF recruits FIGNL1 to damage sites, the reduction in FIGNL1 levels upon RADIF loss and reduced FIGNL1 expression upon cisplatin treatment prevented us from directly determining this. Attempts at addressing this using a FIGNL1-GFP were also as of yet unsuccessful. RADIF binding to FIGNL1 similarly required the N-terminal region, and we demonstrated that mutants of RADIF that failed to bind to FIGNL1 could not rescue the sensitivity of ΔRADIF cells to cisplatin treatment.

Our current model for how RADIF regulates RAD51 is through stabilization and perhaps recruitment of FIGNL1, which performs the effector function of controlling RAD51 levels on chromatin. Purified FIGNL1 promotes dissociation of RAD51 from ssDNA (*44*), providing a biochemical basis for our observed increased persistence of RAD51 in the absence of RADIF-FIGNL1 upon ICL treatment, and the dissociation of RAD51 from foci when RADIF is over-expressed. Our experiments showing striking sensitivity to ICL agents following RADIF-FIGNL1 loss but reduced sensitivity to HU or IR treatment is similar to what was seen with other RAD51 regulatory helicases ZGRF1 and HELQ (*20, 53–55*). These proteins, when absent, show defective HR using reporter assays and lead to increased persistence of RAD51 foci upon DNA damage. Although these all have HR functions, ICL repair appears to be particularly dependent on not just RAD51’s HR role but other functions linked to fork reversal and fork protection. In addition to RADIF, FIGNL1 also interacts with SPIDR (*27*) and SWSAP1 (*44*). SWS1, SWSAP1 and SPIDR together form a distinct complex, that was recently shown to be dispensable for intra-chromosomal HR but essential for inter-homolog HR (*56*), showing that various types of recombination might require multiple different regulators of RAD51 function. How RADIF regulates FIGNL1’s multiple interactions will be a subject of future studies.

RADIF limits RAD51 levels on DNA even in the absence of DNA damage. A similar function has been attributed to RADX, which regulates the amounts of RAD51 at stalled forks (*57–59*) by competing with RAD51 for binding to ssDNA. Unlike RADX, RADIF and FIGNL1 are both required for proper HR consistent with a role downstream of RAD51 foci formation. Too much RAD51 on chromatin may be detrimental to replication as it might be necessary to finetune the fork reversal activities of RAD51 to prevent excessive and/or unwarranted fork reversal (*60*). Failure to do so might lead to increased fork collapse. Consistent with this, we observed increased γH2AX upon RADIF loss. RAD51 can bind to both ssDNA and dsDNA, and its dsDNA binding activity has recently been proposed to be the basis for its role in fork protection (*26*). Whether the observed increased chromatin association of RAD51 upon RADIF loss occurs specifically on nascent ssDNA vs dsDNA or whether FIGNL1 can also dissociate RAD51 will be interesting to discover.

Strikingly, there was a marked reduction in FIGNL1 protein levels upon depletion of RADIF. This was seen following multiple siRNA knockdowns and in both knockout clones. Interestingly, loss of FIGNL1 also led to loss of RADIF expression, revealing that both proteins form a co-stable complex. In addition, both proteins appeared to be regulated differently upon damage, as FIGNL1 but not RADIF was targeted for degradation after cisplatin treatment in a proteasome-dependent manner. FIGNL1’s loss upon DNA damage is reminiscent of other factors like EXO1 (*61, 62*) and CHK1 (*63*), but how and why FIGNL1 is degraded after ICL treatment is not yet clear. One possibility is that the during replication cells maintain just enough RAD51 on DNA to respond to sporadic lesions that arise without compromising normal replicative potential, hence requiring high amounts of FIGNL1 on DNA. However, upon DNA damage when several forks stall and replication slows considerably, FIGNL1 levels on DNA could be reduced to allow increased RAD51 levels to drive its multiple repair functions. How this is coordinated with RAD51 mediators will be of interest. Indeed, whereas FIGNL1 can counteract SWSAP1 activity in regulating RAD51 foci formation, it has no effect on another paralog, RAD51C (*44*). That notwithstanding, complete loss of RADIF-FIGNL1 is problematic as well, as both proteins are likely necessary for dissociating RAD51 from DNA once strand invasion is complete and/or to fine-tune RAD51 nucleofilament formation during homology search to prevent unwanted recombination, hence, the severe sensitivity of RADIF-FIGNL1 null cells to agents such as cisplatin and MMC.

Platinum based chemotherapeutics are widely used in the treatment of several cancer types. We show here that the RADIF-FIGNL1 complex is critical for resistance to chemotherapeutic agents, suggesting that targeting this complex might be a potent combination approach to treatment of several cancers. PARP inhibitors have also recently been widely used for treatment of HR-deficient cancers (*50, 64*). We found that RADIF knockdown potentiates sensitivity to Olaparib treatment, likely because of RADIF-FIGNL1 roles in HR control. The ATPase dead mutant of FIGNL1 is still able to dissociate RAD51 from ssDNA in vitro and may not be an effective therapeutic approach (*44*), however, disruption of the interaction between RADIF and FIGNL1 using small peptides might be a potent strategy for treatment of various cancers.

## ACKNOWLEDGEMENTS

We thank all the members of the Adeyemi Lab for helpful discussions.

## AUTHOR CONTRIBUTIONS

Conceptualization: ROA, Methodology: JDT, HT, ROA, Investigation: JDT, HT, TTO, AP, ROA, Visualization: JDT, HT, ROA, Supervision: ROA, Writing – original draft: ROA, Writing – review & editing: JDT, HT, ROA

## DATA AND MATERIALS AVAILABILITY

All data needed to evaluate the conclusions in the paper are present in the paper and/or the Supplementary Materials.

## MATERIALS AND METHODS

### Cell lines

All U2OS cells were passaged in McCoys 5A media supplemented with 10% fetal bovine serum and 1% penicillin/streptomycin. RPE1 cells were maintained in DMEM:F12 media supplemented with 10% FBS and 1% penicillin/streptomycin. HeLa cells and 293T cells were grown in DMEM supplemented with 10% fetal bovine serum and 1% penicillin/streptomycin. SUM149 cells were grown in Ham’s F12 media supplemented with 10% FBS and 1% penicillin/streptomycin. DR-GFP U2OS cells were previously described (*65*). All cells were maintained at 37°C in 5% CO2, and passaged using 0.25% Trypsin-EDTA to dissociate cells.

### Plasmids and Cloning

Myc-DDK-tagged lenti ORF clone of c1orf112 (Lenti-Myc-FLAG-RADIF) was obtained from Origene (RC211444L1). Human FIGNL1 sequence-verified cDNA (FIGNL1-cDNA) was obtained from Horizon Discovery (MHS6278-202759761). pcDNA3.1-3xFLAG was generated by inserting a DNA fragment coding 3xFLAG tag into the multiple cloning site of pcDNA3.1 using restriction enzymes Xba1 and Apa1. To make 3XFlag-RADIF or 3XFLAG-FIGNL1, each gene was amplified by PCR and cloned into PCDNA3.1-3XFLAG using KpnI and XhoI restriction enzymes sites. N-terminal HA-tagged RAD51 was generated by inserting an HA-RAD51 DNA fragment synthetized as a gBlock fragment by IDT into pcDNA3.1 vector using BamHI and ApaI sites. The RADIF N and C terminal truncations and FIGNL1 truncations were all obtained by PCR cloning. GFP-tagged-RADIF was obtained by restriction digestion from Lenti-Myc-FLAG-RADIF using AsiSI and MluI and cloning into a pCMV6-AC-GFP vector originally obtained from Origene. gRNA-target-resistant constructs were generated by site directed mutagenesis using QuikChange II Site-directed mutagenesis kit (200521, Agilent) or Q5 Site-directed mutagenesis (E0554, NEB) according to the manufacturer’s instructions. All primers used in this study are provided in Supplemental Table 2. All constructs used in this study were verified either by Sanger sequencing or whole plasmid sequencing by Primordium labs.

### Plasmid transfections and virus production/viral transduction

Plasmids were transfected into cells 293T cells using PolyJet (SL100688, Fisher) according to the manufacturer’s instructions. Cells were left untreated or treated with drugs and harvested 2-4 days later. To make stable cell lines, restriction-digest linearized plasmids were transfected into U2OS or HeLa cells using Lipofectamine 3000 reagent according to the manufacturer’s instructions. 24 h later cells were selected using Geneticin for 5-7 days. With GFP-tagged constructs, selected cells were subsequently sorted using BD FACSymphony S6 (BD Biosciences). Lentiviruses were generated as previously described (*28*) in 293T cells, filtered and used to infect target cells.

### siRNA transfections

RPE1, U2OS, 293T or HeLa cells were reverse transfected into cells at 20-40 nM using Lipofectamine RNAiMAX reagent (13-778-075, Invitrogen) according to the manufacturer’s instructions. Occasionally, seeded cells were transfected a second time with siRNAs the next day. Cells were then processed as described and harvested 2-4 days later. Unless otherwise indicated siRNAs used in this study were SmartPools from Dharmacon. Control siRNAs used were Allstars Negative control siRNAs (Qiagen).

### Generation of CRISPR Knockout RADIF (c1orf112) cells

gRNA sequences were annealed and cloned into LentiCrispr v2 (Addgene) modified to express NAT resistance gene. After virus generation, U2OS cells were infected and selected using 250 µg/mL of NAT for 5 days. Single cells were cloned and RADIF knockout status was ascertained by western blotting. To confirm knockout status, genomic DNA was isolated using GeneJet Genomic DNA Purification Kit (K0702, ThermoFisher Scientific). gRNA target knockout regions were amplified by PCR and Sanger sequenced.

### Reagents

DMSO was obtained from Fisher Scientific (Cat. # 97063-136). Cisplatin was purchased from Selleck Chemicals (Cat. # S1166). Hydroxyurea was obtained from Sigma (Cat. # H8627) and dissolved in PBS. CD437 was obtained from Tocris Bioscience (Cat. # 1549). Olaparib was from Selleck Chemicals (Cat. # S1060). MMC was obtained from Santa Cruz Biotechnology (Cat. # sc-3514A). Puromycin from Sigma (Cat. # P8833). Geneticin from Gibco (Cat. # 0131035). Nourseothricin (NAT) Gold Biotechnology (Cat. # 501532818). Halt Protease and Phosphatase Inhibitor Cocktail was obtained from Fisher Biotech (Cat. # 78440).

### Immunoblots

Western blotting was done as previously described (*28*). Briefly, cells were harvested and lysed in RIPA buffer on ice for 15 – 30 minutes, clarified and transferred to new tubes. Alternatively, for whole cell lysates, cells were harvested and lysed in SDS lysis buffer (50 Mm Tris HCl pH 6.8, 100 mM NaCl, 1% SDS, 10 mM NaF, 7% glycerol) prior to sonication. Protein content was measured on an Eppendorf Biophotometer using Bradford reagent, sample buffer was added, and samples were analyzed using sodium dodecyl sulfate gel electrophoresis (SDS-PAGE). Membranes were blocked in 5% (wt/vol) milk in Tris-buffered saline with Tween (TBST) buffer and then probed with antibodies (see antibodies). Westerns were quantified using ImageJ.

### Antibodies

The antibodies used in this work are as follows: Anti-HA (Sigma, H3663-200), c1orf112 (Sigma, HPA023778), FANCD2 (Santa Cruz Biotechnology, D0114), FANCA (Bethyl, A301-980A), FANCI (Bethyl, A301-254A), Vinculin (Sigma, V9131-.2ML), ATM (Abcam, ab81292), γH2AX (Sigma, 2884537), mouse monoclonal anti-FLAG (Sigma, F1804-200UG), Rabbit anti-GFP antibody (Abcam, ab6556), RPA32-P-S4/8 (Bethyl, A300-245A), RPA32 (Santa Cruz Biotechnology, F0420), CHK1-P-S317 (Cell Signaling, 2344S), CHK1 (Santa Cruz Biotechnology, I2515), FIGNL1 (Proteintech, 17604-1-AP-150UL), GAPDH (Santa Cruz Biotechnology, sc-47724), ORC2 (Abcam, ab68348), RAD51 (Abcam, ab63801), Mouse monoclonal RAD51 (Millipore-Sigma, Q2574784), Rad51 Antibody (H-92) (Santa Cruz Biotechnology, sc-8349), Goat anti-Rabbit IgG (H+L) Highly Cross-Adsorbed Secondary Antibody, Alexa Fluor Plus 488 (Invitrogen, A32731), Goat anti-Mouse IgG (H+L) Highly Cross-Adsorbed Secondary Antibody, Alexa Fluor Plus 488 (Invitrogen, A32723), Goat anti-Rat IgG (H+L) Highly Cross-Adsorbed Secondary Antibody, Alexa Fluor Plus 488 (Invitrogen, A48262), Goat anti-Mouse IgG (H+L) Highly Cross-Adsorbed Secondary Antibody, Alexa Fluor Plus 594 (Invitrogen, A32742), Goat anti-Rabbit IgG (H+L) Highly Cross-Adsorbed Secondary Antibody, Alexa Fluor Plus 594 (Invitrogen, A32740).

### Co-IP analyses

293T cells were transfected with the indicated plasmids for 48 – 60 hours in 60 mm dishes. Occasionally, samples were treated with cisplatin or vehicle for 18-24 hours. Cells were harvested in cold PBS and lysed in RIPA buffer without SDS. Lysates were clarified by centrifugation and the supernatants were precleared with Pierce™ Protein A/G Magnetic Beads (88802, Thermofisher) and immunoprecipitated using Anti-FLAG® M2 Magnetic Beads (M8823, Sigma) for 4 hours at 4°C. Beads were washed three times and boiled in sample buffer prior to western blotting.

### Chromatin fractionation

WT and/or RADIF KO U2OS cells previously transfected or not with the indicated siRNAs were seeded in 10cm plates, treated with drugs as indicated, harvested and washed with cold PBS. Sedimented cells were resuspended in cold Solution 1 consisting of 10 mM Hepes (pH 7.9), 10 mM KCl, 1.5 mM MgCl_2_, 0.34 M sucrose, 1 mM DTT, protease and phosphatase inhibitor cocktails. Triton X-100 was added to 0.1%. After a 5-minute incubation, samples were centrifuged at 1300g for 5 minutes, and the supernatant removed as the soluble fraction. Sedimented nuclei were washed once with Solution 1 and lysed in Solution 2 (3 mM EDTA, 0.2 mM EGTA, 1 mM DTT, protease and phosphatase inhibitor cocktails) for 30 minutes on ice. Samples were centrifuged at 1300 g for 5 minutes, and the chromatin-enriched pellets washed once with Solution 2 followed by resuspension in lysis buffer containing 50mM Tris pH 6.8, 100 mM NaCl, 1.7% SDS, 7% glycerol and protease and phosphatase inhibitors. After sonication, protein content was measured. Sample buffer was added to 1X and the samples were boiled for western blotting.

### Immunofluorescence Assays

U2OS or HeLa cells were transfected or not with the indicated siRNAs, then 24 hrs later plated onto glass coverslips in 6-well plates. After 24–36 hrs, drug treatments were performed for the indicated durations. Cells were then washed with PBS, fixed with 4% paraformaldehyde for 15 min and extracted with 0.5% Triton X-100 in PBS for 10 min. Alternatively, cells were pre-extracted using cytoskeleton buffer (containing 10 mM piperazine-N,N′-bis(2-ethanesulfonic acid) (PIPES), pH 6.8, 100 mM NaCl, 300 mM sucrose, 1 mM MgCl_2_, 1 mM EGTA as well as 0.5% Triton X-100, protease and phosphatase inhibitors) for 5 mins on ice prior to the fixation and permeabilization steps. Cells were then blocked in 3% BSA in PBS and incubated with primary and secondary antibodies. The coverslips were mounted, and nuclei were visualized with DAPI Fluoromount-G (Southern Biotech, OB010020) and images were acquired using an LSM 780 NLO (Zeiss).

### Multicolor competition assay (MCA)

GFP-labeled U2OS cells were reverse transfected with the indicated siRNAs at 20 nM using Lipofectamine RNAiMAX reagent (Invitrogen) while RFP-labeled U2OS cells were transfected with control siRNAs in the same way. The following day, transfection was repeated with the same siRNAs and transfection reagent to maximize the knockdown effect. GFP- and RFP-labeled cells were mixed in equal quantities in six-well plates 2 days after the first siRNA transfection, and they were treated with the indicated dose of drug or vehicle control for 24 h. Fresh media was added and cells were maintained for 6 days after treatment. Subsequently, the percentage of GFP and RFP labeled cells were quantified by FACS analyses using BD FACS Canto II. Data were analyzed using FlowJo software.

### Cell cycle analyses

Cell cycle analyses were performed as previously reported (*28*). Briefly, siRNA treated WT U2OS and/or RADIF knockout cells were treated or not with drugs as indicated. Cells were then harvested, fixed in 4% paraformaldehyde for 15 min at room temperature, or cells were fixed in 70% ethanol for 15 min on ice. After pelleting, cells were washed in PBS, resuspended in 50 µg/ml propidium iodide solution containing 0.1 mg/ml RNase A as well as 0.05% Trition X-100 for 40 min at 37°C, resuspended in PBS and flow cytometry was performed using BD FACSymphony A5 or BD FACS Canto II. Data were analyzed using Flowjo software.

### DNA fiber assay

U2OS cells were incubated with 25 µM CldU for 30 min, washed and subsequently treated with 250 µM IdU for 30 min. After labeling, cells were washed, harvested and resuspended in PBS. 2 μL of the cell suspension were transferred to a glass microscope slide, overlaid with 6 μL lysis buffer (0.5% SDS, 200 mM Tris-HCl (pH 7.4), and 50 mM EDTA), and the slide was tilted to allow DNA to spread by gravity. After air-drying, 3:1 methanol/acetic acid was applied on the slides to fix DNA. DNA was denatured by incubating the slide in 2.5 M HCl for 80 min, followed by wash with PBS. Blocking was performed with 5% BSA in PBS for 30 min. For immunostaining, slides were incubated overnight with primary antibodies; ab6326 anti-BrdU (cross-reacts with CldU) antibody (rat) (1:100) and BD Biosciences 347580 anti-BrdU (cross-reacts with ldU) antibody (mouse) (1:25). Slides were washed with PBS followed by incubation for one hour with the secondary antibodies; anti-rat AIexa-488 antibody (1:400) and anti-mouse Alexa-594 antibody (1:400). After wash with PBS, mounting medium was added on the slides and images were acquired with Leica SP8 confocal microscope. Images were analyzed with ImageJ. More than 200 fibers were counted for each condition.

### Metaphase spreads

HeLa cells were reverse transfected with the indicated siRNAs at 20 nM using Lipofectamine RNAiMAX reagent (Invitrogen). The following day, transfection was repeated with the same siRNAs and transfection reagent to maximize the knockdown effect. Two days after from the first transfection, cells were treated with 2 ng/ml MMC or PBS for 48 h. Following treatment, the cells were exposed to colcemid (100 ng/ml) for 2 h, treated with a hypotonic solution (1:2 mixture of 0.075 M KCl and 0.9% Na citrate) for 10 min and fixed with 3:1 methanol: acetic acid. Slides were stained with Giemsa stain and 50 metaphase spreads were scored for aberrations. Metaphase spreads were observed using a Zeiss Axiovert 200M microscope and captured using AxioVision.

### Clonogenic survival assay

Colony formation assays were performed as previously described (*28*). Briefly, where indicated, cells were reverse transfected with siRNA to the indicated genes as described above. Afterwards, WT or CRISPR KO cells were exposed to the indicated doses of drugs for 16 to 24 h and adjusted for plating depending on dose of drug in six-well plates. After 7-10 days, cells were fixed and stained using crystal violet and scored with a colony counting pen (VWR).

### Cell viability assays

Cell viability assays using CellTiter-Glo 2 (Promega) were performed according to the manufacturers’ instructions. Briefly, following siRNA treatment, cells were seeded at 500 to 1000 cells per well in triplicate in 96 well plates. Cells were then treated with the indicated drugs for 24 hours, washed and left to recover for 48 to 72 hours prior to being read on a BioTek FLx800.

### Homologous Recombination assay

U2OS cells with a stably integrated DR-GFP reporter (*66*) were transfected with the indicated siRNAs 48 h prior to infection with adenovirus expressing I-SceI restriction enzyme at an MOI of 10. The percentage of GFP-positive cells was determined 48 h after infection by flow cytometry using a BD FACSymphony A5 (Becton Dickinson). Data were collected using BD FACS Diva software (Becton Dickinson), and analysis was performed using FlowJo Software.

### Quantification and statistical analyses

Statistical analyses were performed using Prism 9 (GraphPad). A Mann-Whitney nonparametric test was used when comparing two samples. An analysis of variance (ANOVA) test was used when comparing more than two groups, and if there was a difference, this was determined by a Dunn’s multiple comparisons post-test. Multiple siRNAs, multiple knockout clones, and multiple cell lines were analyzed to confirm that results were not caused by off-target effects or clonal variations. For all experiments: n.s. P ≥ 0.05; *P < 0.05; **P < 0.01; ***P < 0.001; ****P<0.0001. All experiments in this work were performed at least twice, and representative experiments are shown.

## SUPPLEMENTAL MATERIALS

**Fig. S1.**
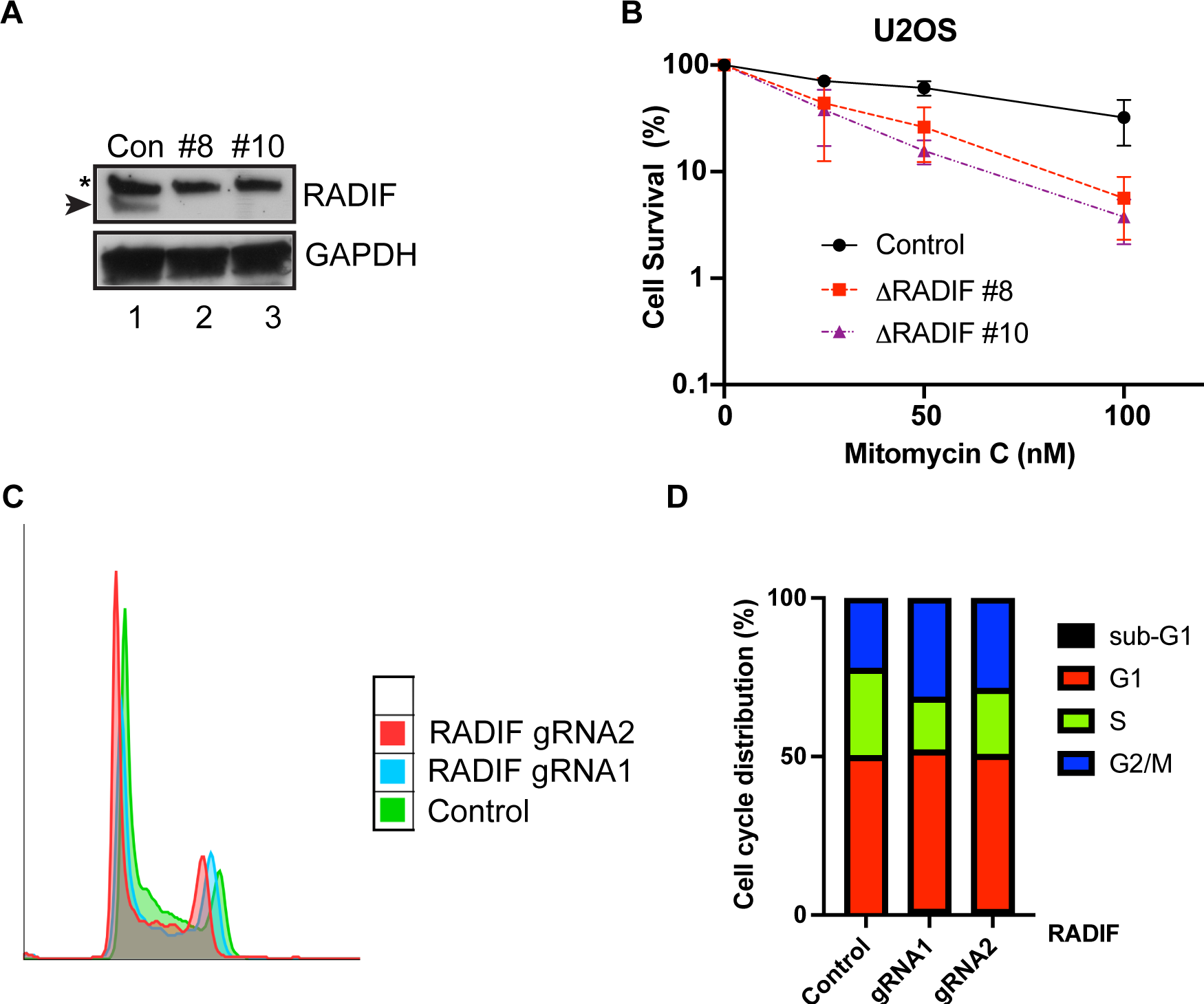
**(A)** Western blots showing loss of RADIF in both RADIF knockout clones (#8 and #10). Con – control gRNA expressing U2OS cells. **(B)** CSAs showing survival of U2OS cells expressing control gRNA or two RADIF gRNA knockout clones upon treatment with indicated doses of MMC. Mean ± SD of two independent experiments are shown. **(C-D)** Representative images of cell cycle distribution in control or RADIF gRNA expressing U2OS cells overlayed in the absence of drug treatment, quantified in (D).

**Fig. S2.**
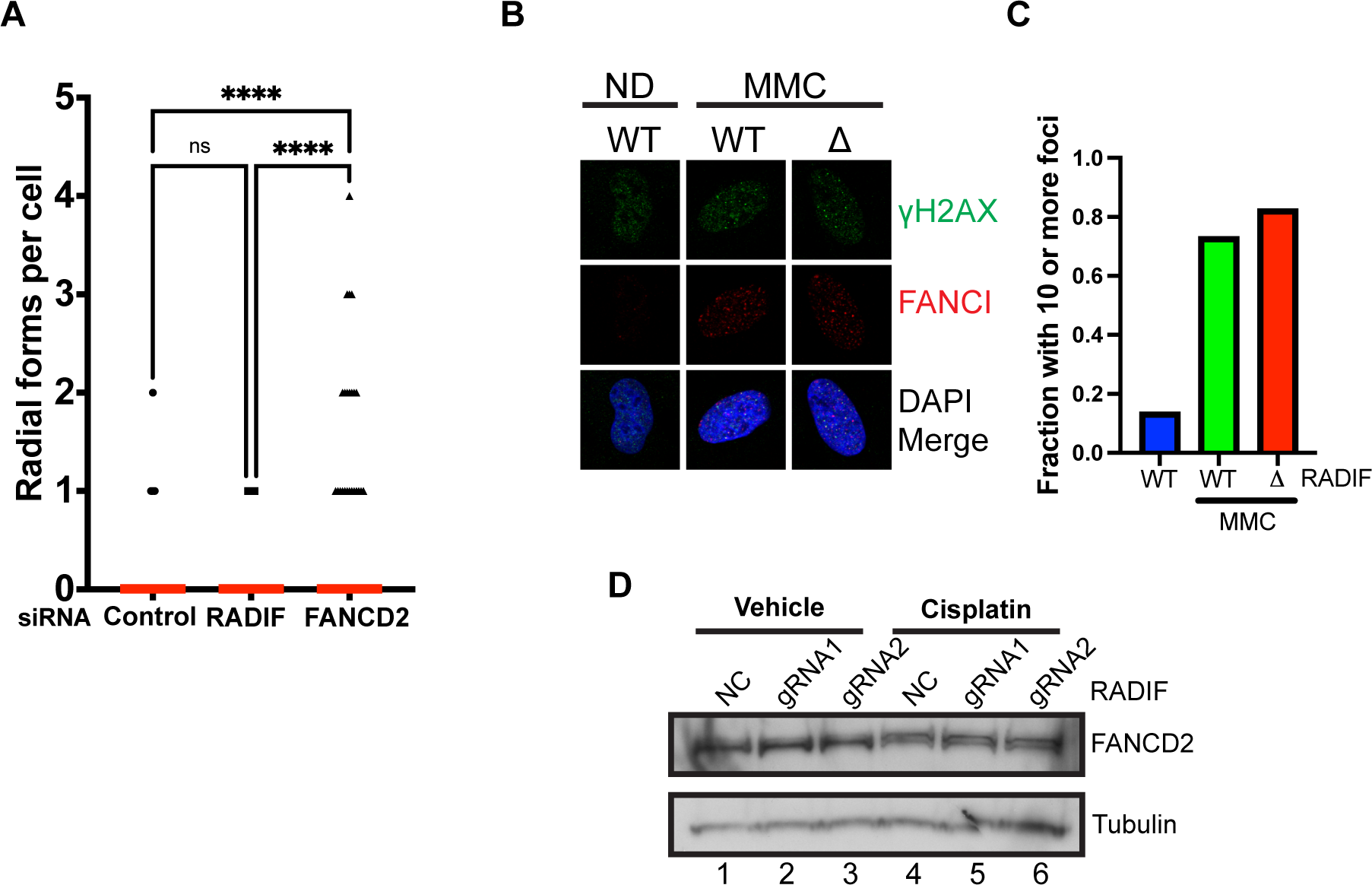
**(A)** Numbers of radial form chromosomes at metaphase in MMC-treated HeLa cells transfected with indicated siRNAs. Red bar is median of each group. **(B)** U2OS cells expressing control or RADIF gRNA were treated with 1 μM MMC for 8 hours prior to IF analyses for FANCI and γH2AX foci. **(C)** Quantification of the experiment in (B). **(D)** Western blot analyses showing unchanged FANCD2 ubiquitination following cisplatin treatment in U2OS cells expressing two different gRNAs to RADIF or control gRNA (NC).

**Fig. S3.**
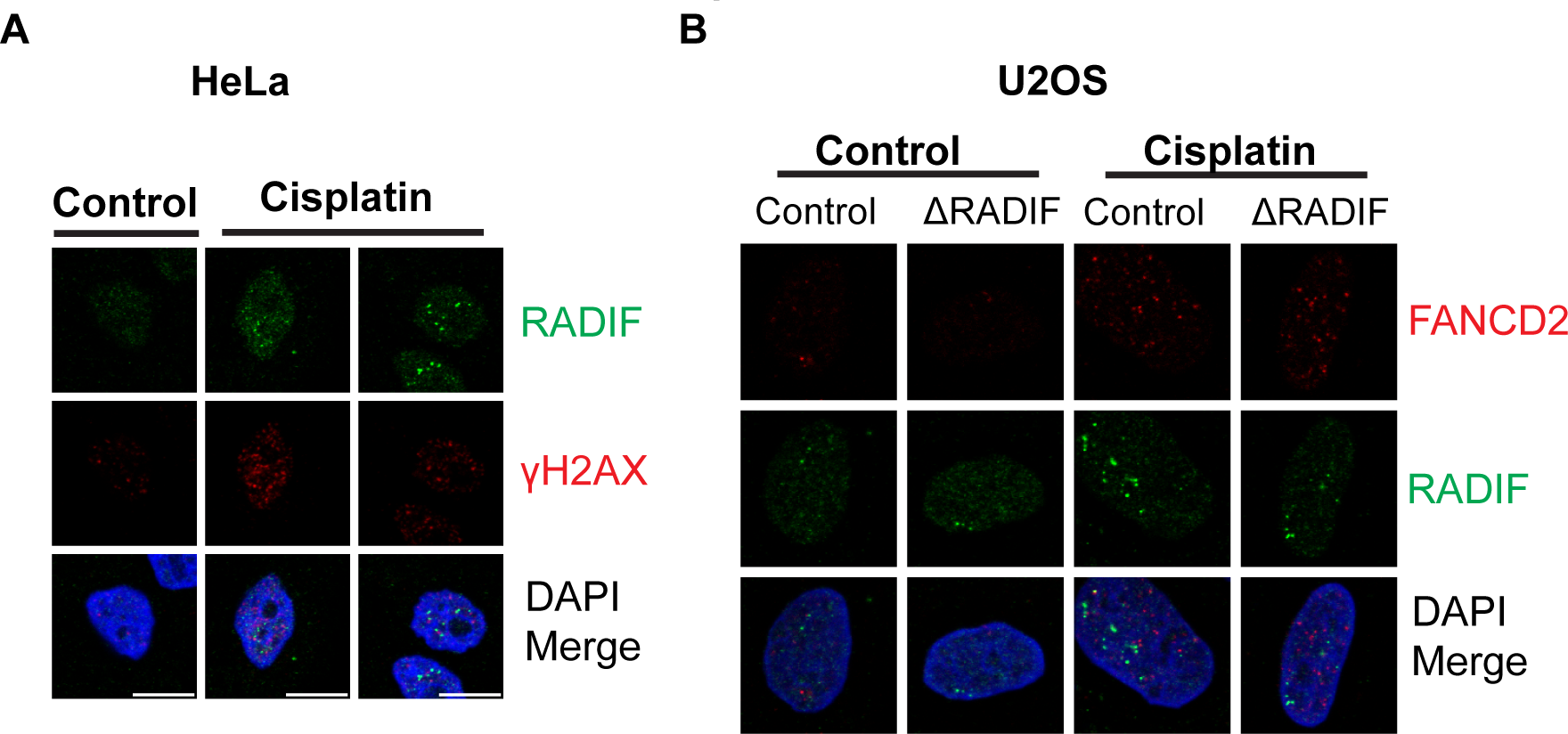
**(A)** IF images showing recruitment of GFP-RADIF to foci in HeLa cells after treatment with 0.5 μM cisplatin for 16 h. Cells were pre-extracted to remove soluble protein. **(B)** IF images in U2OS cells showing recruitment of GFP-RADIF to foci that only partly colocalize with FANCD2 after treatment with 0.5 μM cisplatin for 16 h. Cells were pre-extracted to remove soluble protein.

**Fig. S4.**
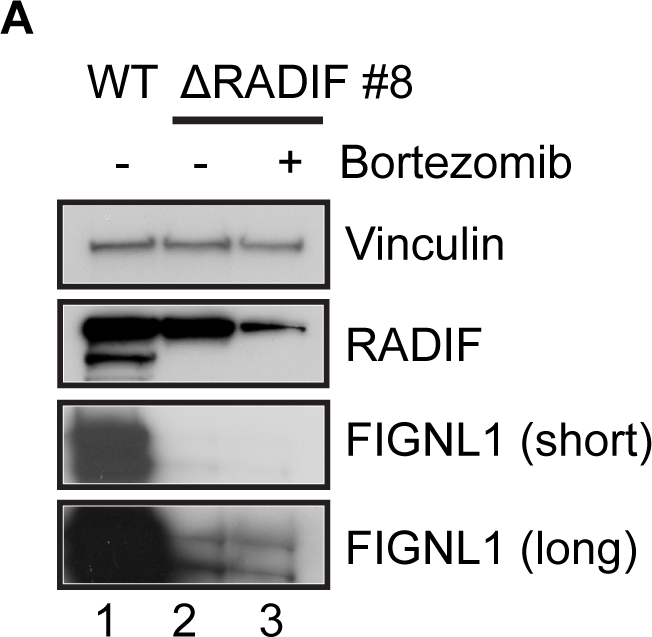
**(A)** Control gRNA expressing U2OS cells (WT) or 1′RADIF knockouts were treated with vehicle or 10 Bortezomib for 14 hrs as indicated. WCLs were prepared and assayed for the indicated proteins by WB.

**Fig. S5.**
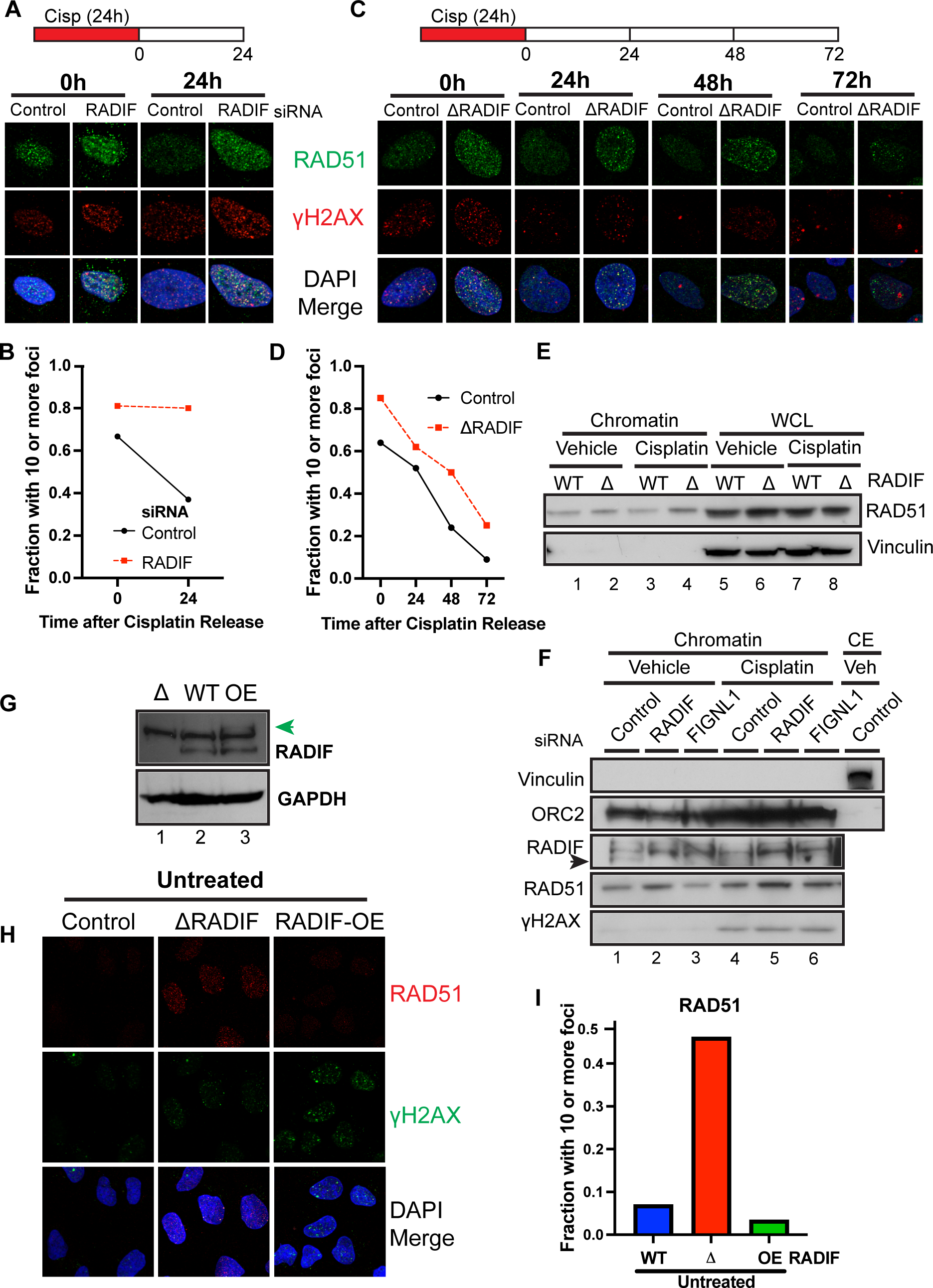
**(A)** Top: Schematic of the experiment. U2OS cells were transfected with the indicated siRNAs for 48 h prior to 1 μM cisplatin treatment as indicated. Bottom: Representative IF images show RAD51 and γH2AX foci at different time points after release from cisplatin treatment. **(B)** Quantification of the experiment in (A). **(C)** Top: Schematic of the experiment. Control gRNA expressing or 1′RADIF U2OS cells were treated with 0.5 μM cisplatin as indicated. Bottom: Representative IF images show RAD51 and γH2AX foci at different time points after release from cisplatin treatment. **(D)** Quantification of the experiment in (C). **(E)** Control gRNA-expressing (WT) or 1′RADIF U2OS cells were treated with vehicle or 1.5 μM cisplatin for 18 h. Samples were split 2:1 ratio then processed for chromatin fraction or WCLs prior to blotting for the indicated proteins. **(F)** U2OS cells were reverse transfected with control, RADIF or FIGNL1 siRNAs for 48h prior to treatment with vehicle or 2.5 μM cisplatin for 18 h. Cells were fractionated and blotted for the indicated proteins. **(G)** Western blots show RADIF levels in knockout (Δ), U2OS (WT) and RADIF-GFP expressing (OE, green arrow) cells. **(H)** Control, 1′RADIF or RADIF-GFP overexpressing (OE) U2OS cells were processed for IF. Representative IF images show RAD51 and γH2AX foci. **(I)** Quantification of (H).

**Fig. S6.**
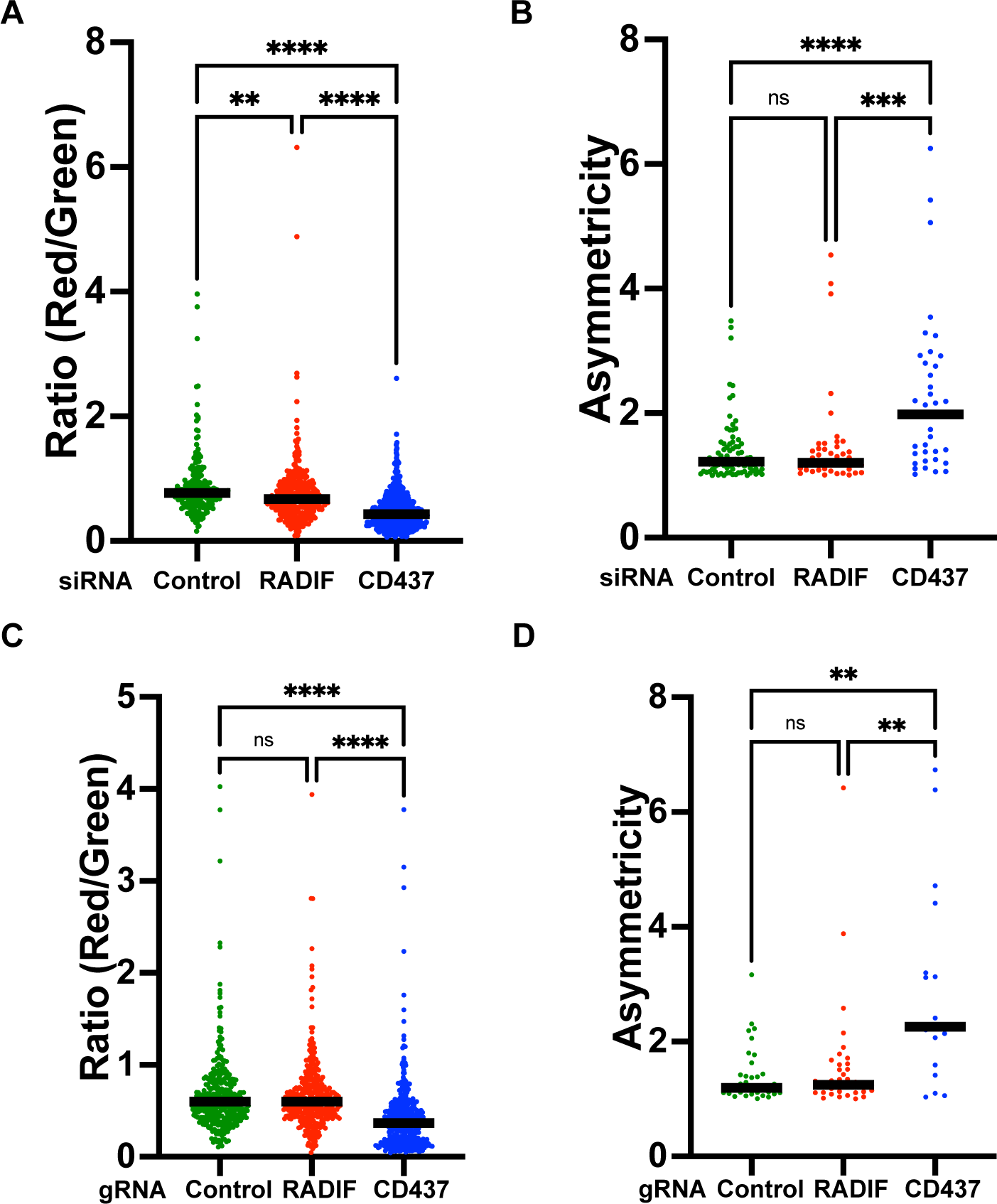
**(A)** Ratios (red/green) of the length of two-color DNA fibers obtained from U2OS cells transfected with siRNAs as indicated or treated with CD437. **(B)** Quantified asymmetricity of the bidirectionally labeled DNA replication fork from DNA fiber assay. The ratios of the length of two red fibers in three-segmented DNA fibers (red-green-red) were calculated (long red/short red). DNA fibers were obtained from U2OS cells transfected with siRNAs as indicated or treated with CD437. **(C)** Same as (A). DNA fibers were obtained from U2OS cells expressing indicated gRNAs, or control gRNA with CD437 treatment. **(D)** Same as (B). DNA fibers were obtained from U2OS cells expressing indicated gRNAs, or control gRNA with CD437 treatment.

**Table S1.**
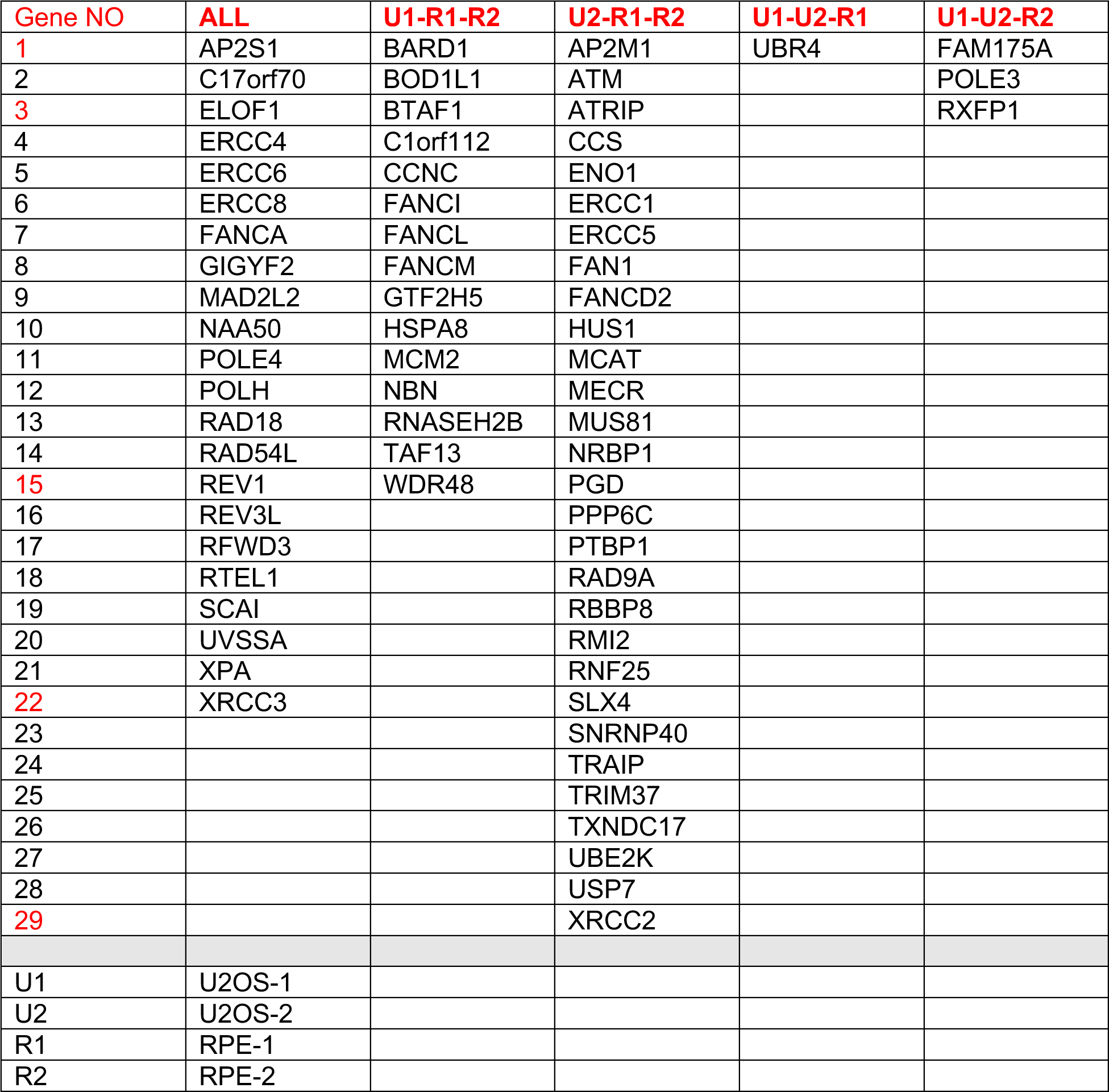
Overlap of top U2OS and RPE genome-wide cisplatin screen hits.

**Table S2.**
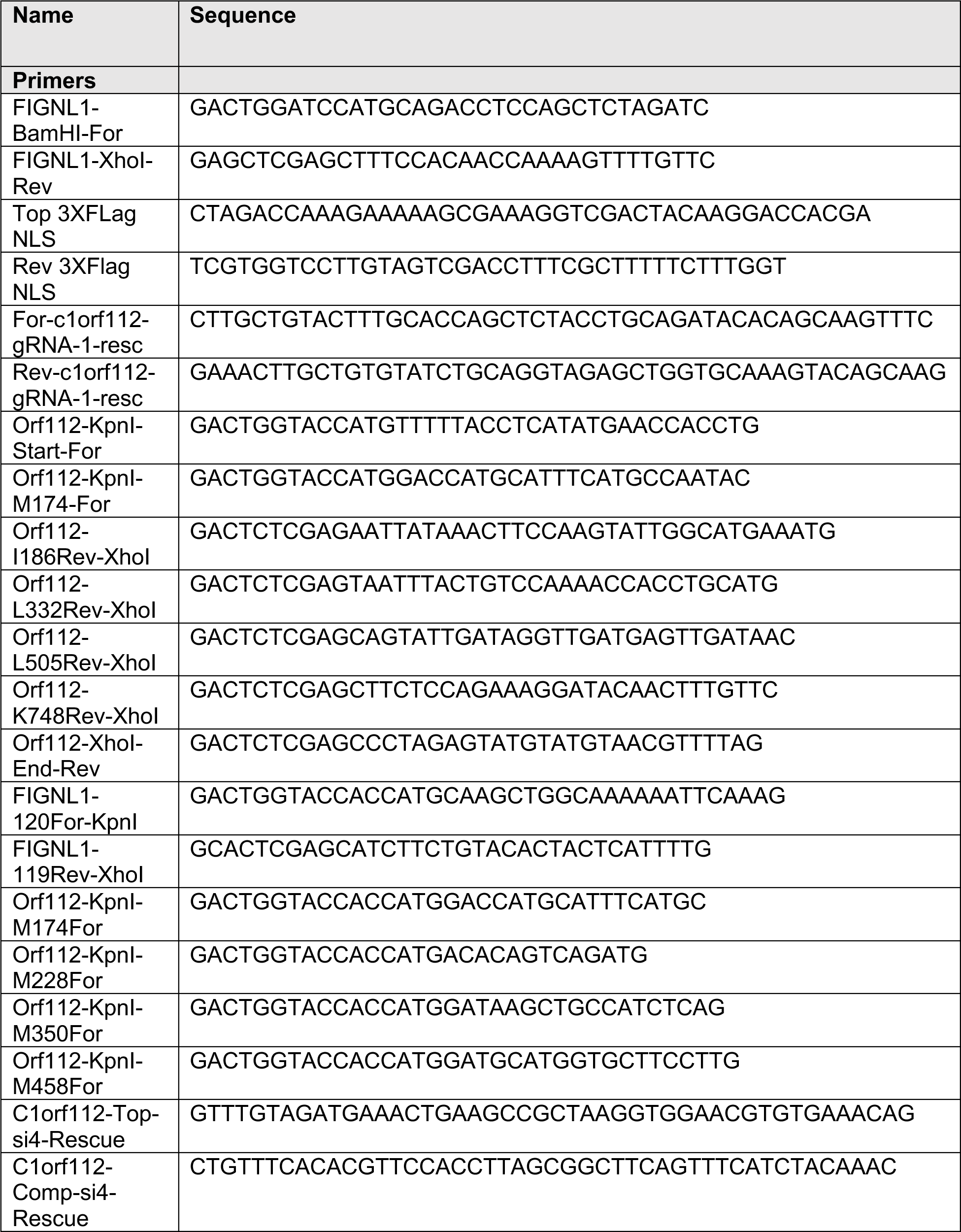
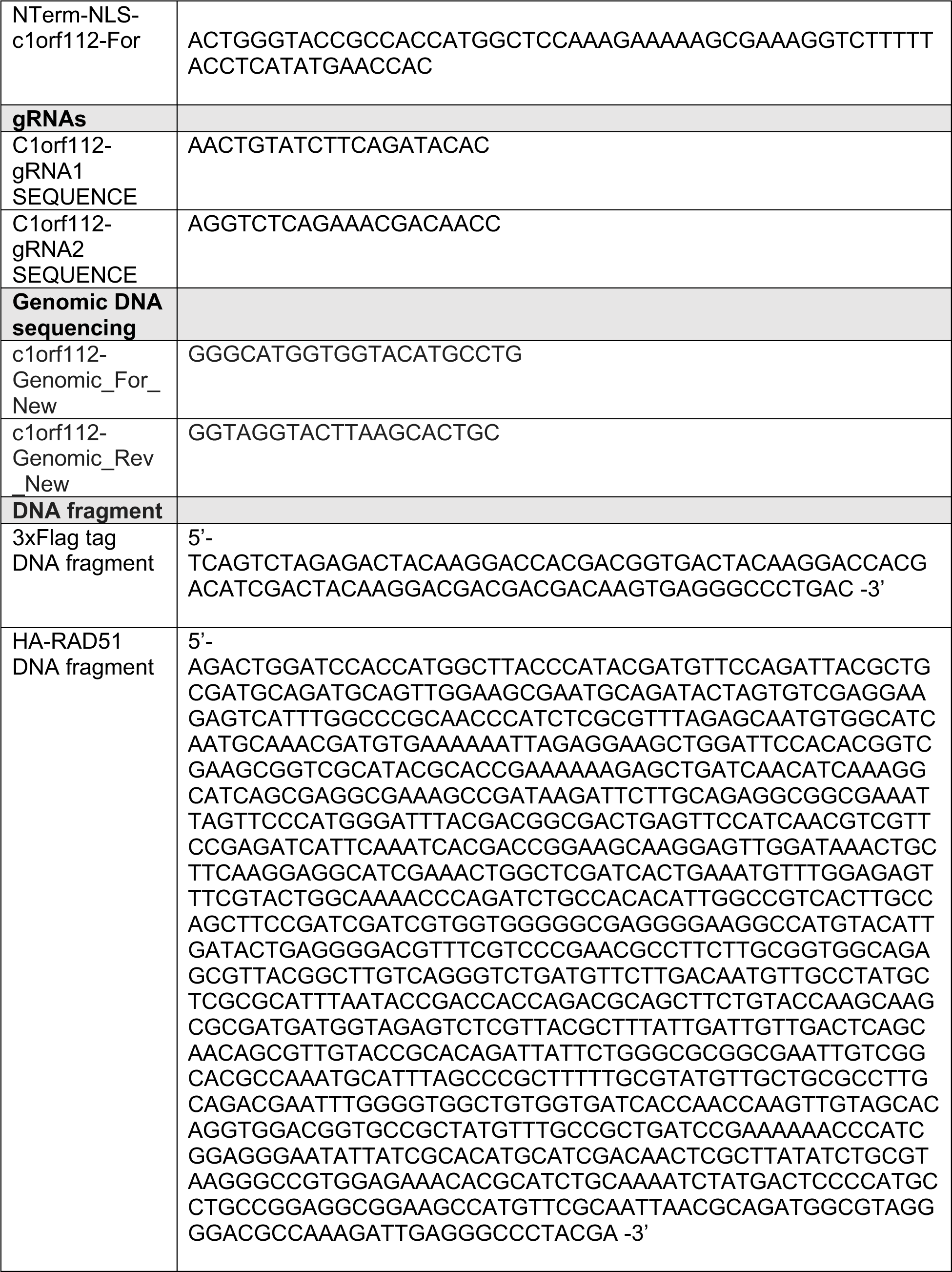
List of primers and gene fragments used.

